# Dietary restriction reveals sex-specific expression of the mTOR pathway genes in Japanese quails

**DOI:** 10.1101/2024.03.10.584307

**Authors:** Gebrehaweria K. Reda, Sawadi F. Ndunguru, Brigitta Csernus, Renáta Knop, James K. Lugata, Csaba Szabó, Levente Czeglédi, Ádám Z. Lendvai

## Abstract

Limited resources affect an organism’s physiology through the conserved metabolic pathway, the mechanistic target of rapamycin (mTOR). Males and females often react differently to nutritional limitation, but whether it leads to differential mTOR pathway expression remains unknown. Recently, we found that dietary restriction (DR) induced significant changes in the expression of mTOR pathway genes in female Japanese quails (*Coturnix japonica*). We simultaneously exposed 32 male and female Japanese quails to either 20%, 30%, 40% restriction or *ad libitum* feeding for 14 days and determined the expression of six key genes of the mTOR pathway in the liver to investigate sex differences in the expression patterns. We found that DR significantly reduced body mass, albeit the effect was milder in males compared to females. We observed sex-specific liver gene expression. DR downregulated *mTOR* expression more in females than in males. Under moderate DR, *ATG9A* and *RPS6K1* were increased more in males than in females. Like females, body mass in males was correlated positively with *mTOR* and *IGF1,* but negatively with *ATG9A* and *RS6K1*. Our findings highlight that sexes may cope with nutritional deficits differently and emphasise the importance of considering sexual differences in studies of dietary restriction.

## Introduction

Nutritional availability is intricately linked to challenges for survival in the natural environment. In natural habitats, animals contend with seasonal fluctuations in food supply and encounter obstacles such as topographical barriers, predation risks, distances, etc., which may expose them to temporary and/or seasonal episodes of starvation. This dynamic interplay with resource availability influences essential life-history traits, including growth, reproduction, and survival^1–3^.

Dietary restriction (DR) is one of the most robust interventions used to study the effect of resource limitation on phenotypic, physiological and molecular plasticity. DR affects life-history traits antagonistically; it improves survival at the expense of growth and current reproduction^4^. In response to nutritional limitations, organisms undergo critical changes in gene expression to adapt their energy metabolism to the prevailing conditions through molecular and physiological functions^5,6^. Assessing resource availability and orchestrating plastic adaptive responses to its changes are controlled by nutrient-sensing pathways^7^.

One of the key signalling pathways monitoring nutrient availability is the mechanistic target of rapamycin (mTOR)^8,9^. This pathway comprehends a series of cross-talking genes at different stages of cellular functioning. mTOR is an intracellular serine/threonine protein kinase that plays a crucial role in protein synthesis, cell growth, differentiation, and subsequent organismal growth and reproduction^10^. In response to growth hormones and energy availability, mTOR receives signal transduction from extracellular growth factors, mainly insulin-like growth factor 1 (IGF-1) binding to its receptor (IGF-1R) at the plasma membrane, which, in turn, activates downstream effectors by phosphorylation^11,12^. Nutrients (intracellular amino acids) also directly regulate mTOR activity^13,14^. The mTOR is then responsible for activating and inhibiting several transcription and translation factors and binding proteins, ultimately affecting gene and protein expression under the pathway^15^.

The majority of studies exploring DR have mainly focused on the effect of DR on protein expression and posttranslational modification in well-established model organisms such as worms, flies, and rodents^16–18^. However, investigating the role of mTOR in birds is particularly important because of their unique physiology. Birds have high metabolic rates and demand substantial energy reserves for flight, reproduction, and maintaining high body temperature^19,20^. Therefore, by investigating the regulation of mTOR pathway in birds, we not only gain insights into the fundamental mechanisms governing the physiological adaptations of this evolutionarily independent lineage but also unlock a rich source of information with broader implications for understanding the evolutionary and ecological dynamics of nutrient sensing in a diverse spectrum of organisms.

Exploring the distinction between male and female phenotypes and genes that show sex-specific expression (sex-linked) has long been the interest of biologists. Recent studies identified somatic genes differentially expressed across different tissues in response to treatment in males and females in mammals and fly models^21–24^. In mammals, theoretical and empirical evidence shows strong sexual differences in the mTOR-mediated life history regulation^25–28^. For instance, in mice, a moderate dietary restriction (20% DR) improves health span for both males and females, while a severe restriction (40% DR) is detrimental for females but still increases lifespan in males due to divergent physiological and molecular responses. In fruit flies, the effects of dietary restriction on lifespan and mortality rates also differ between sexes, with females showing a peak in lifespan at higher food concentrations and a more pronounced response to restriction^29^. Another experiment in the same species revealed that dietary restriction-mediated sex differences in fitness are associated with sex-specific effects on the expression of genes mediating the mTOR pathway^21,22^. The proposed explanation for these differences is rooted in sexual variations in nutrient requirements and energy allocation. Divergent reproductive strategies, the modulating function of sex hormones and specific optimal diets for reproduction are among the suggested reasons for the difference in the expression of genes in males and females^1,30^. Because of these inherent physiological and reproductive differences, the response to dietary restriction could be sex-specific^20,25,31^. However, despite the theoretical and empirical evidence in other taxa, the sex differences in mTOR pathway response to DR in birds remain unexplored.

Recently, we have shown that liver genes governing the mTOR signalling pathway expressed differentially across dietary restriction gradients and were related to patterns of changes in body mass and reproductive parameters in female Japanese quails (*Coturnix japonica*)^32^. Specifically, DR downregulated the expression of liver *mTOR*, *IGF1* (insulin-like growth factor 1) and its receptor (*IGF1R*), whereas genes downstream to mTOR, such as ribosomal protein kinase 1 *(RPS6K1)* and autophagy-related 9A (*ATG9A),* showed an increasing trend with the level of restriction. However, males and females may differ markedly in their life history and physiology. This study is therefore performed to test whether the liver gene expression signatures and the corresponding fitness traits of male quails are consistent with those observed in females. Japanese quails are sexually size-dimorphic, with females being larger and having a more intensive reproductive investment than males. Therefore, we predicted that the sex-specific size difference would correspond to variations in the expression of genes governing the mTOR pathway in the liver in response to experimental manipulation of food availability. We targeted the hepatic gene expression as the liver plays a central role in the complex metabolic pathway of nutrients. Genes involved in nutrient sensing pathways show distinct expression patterns in the liver and are strongly associated with the body’s overall functioning, ultimately influencing fitness-related traits^33^.

## Material and Methods

### Experimental animal management

Four weeks old Japanese quail (*Coturnix japonica*) chicks, containing both sexes, were obtained from a commercial quail breeder (Budai Fürjészet, Hungary) and housed in the Animal House of the Institute of Animal Science, Biotechnology and Nature Conservation of the University of Debrecen (Hungary). Birds were maintained in an experimental house until they reached maturity for an additional 4 weeks before being subjected to the acclimation of experimental conditions. At the age of 8 weeks, 32 male birds with similar body mass were selected for acclimation and assigned to individual cages alongside another 32 female birds from the same batch and reared in the same housing condition ^32^. We kept them under *ad libitum* feed and water for 7 days of acclimation for individual living and the experimental room conditions. The experiment room was maintained under 24 ± 3 °C temperature, 60% - 75% relative humidity and 12:12 h Light:Dark daily photoperiod cycle. The basal feed for experimental quails was formulated as a breeder quail ration (20% crude protein; 12.13 MJ/kg metabolisable energy) based on corn, soybean, and wheat (Table <u>S1</u>)^34^.

### Experimental design

Before the beginning of the experimental treatment, we measured daily feed offered and leftovers of individual birds for 7 days to estimate their daily feed intake. Feed was offered in a 200 g capacity plastic feeder designed to avoid feed spillage. We also measured the live body mass of each bird at the beginning and at the end of the acclimation period to analyse mass change. We aimed to start the experimental treatment once the birds had stopped growing. At the age of 9 weeks, where the experiment started, male and female birds were randomly allocated to four dietary treatments. The birds in each treatment group were provided with 80% (DR20), 70% (DR30), and 60% (DR40) of their average individual feed intake, while the control group was fed ad libitum (ADL). Males and females were kept in the same room and received identical dietary treatments. The average daily feed intake for the ADL, DR20, DR30 and DR40 during the acclimation period is reported in Supplementary Table S2 online. To control for any potential slight environmental variation of the cages (e.g. due to differences in light intensity), we divided the cages into eight blocks based on their vertical position in the cage system’s staircase, and allocated the birds into these blocks. Each block consisted of an equal number of males and females from each treatment group. The experiment, for both male and female groups, was conducted for 14 days on the same condition. Daily feed left in the ADL group was measured and analysed to monitor any significant change in temporal intake, but we found none.

### Measurements and sampling

We measured body mass at the beginning of the experiment (day 0) and on day 7 and 14 using a digital balance (± 0.1 g). On day 14 of the experiment, all birds were euthanised by cervical dislocation by professional veterinarians after sedation with midazolam (5 mg/mL, EGIS Pharmaceuticals PLC, Hungary) and immediately dissected for liver tissue sampling, starting 8:00 am in the morning. To minimise the short-term impact of feeding, we conducted measurements and sampling on an empty gut. To achieve this, we removed the feeders from all birds early in the morning (8:00 am) before the automatic lights were turned on. The collected liver tissue samples were placed in a collection tube, rapidly frozen on dry ice, immediately taken to the laboratory, and stored at −80LJ until further assays. Male and female samples were collected at the same time and were handled in an identical way^32^.

### RNA Extraction and the Real-time quantitative PCR (qPCR)

Total RNA was isolated from the liver tissue using the TRIzol reagent, following the manufacturer’s protocol, which included DNase treatment to prevent DNA contamination (Direct-zol™ RNA MiniPrep, Zymo Research Corporation, U.S). Briefly, 25 - 30 mg of sample was lysed in 600 µl of TRIzol reagent using a D1000 handheld homogeniser (Benchmark Scientific, USA), and then centrifuged at 16,000 g for 30 seconds at 4 °C to remove debris. The supernatant was then transferred into an RNase-free tube and thoroughly mixed with an equal amount of ethanol (95-100%). The mixture was then transferred into a Zymo-Spin™ IIC column on a collection tube and centrifuged at 16,000 g for 30 seconds. The flow-through was discarded, and the RNA pellet was washed using 400 µl of RNA wash buffer, repeating the centrifugation step. Next, a DNA digestion step was performed by adding 5 µl of DNase I (6 U/µl) and 75 µl of DNA digestion buffer and incubating at room temperature for 15 minutes to purify the RNA from DNA. After adding 400 µl of Direct-zol™ RNA PreWash, we centrifuged for 30 seconds and repeated this step. For the final wash, we added 700 µl of RNA wash buffer and centrifuged for 2 minutes to ensure the complete removal of the wash buffer. Finally, we collected the purified RNA by adding 50 µl of DNase/RNase-Free water for further RNA quality and quantity check, and cDNA synthesis. The RNA concentration and purity were assessed using the HTX Synergy Multi-Mode Microplate Reader spectrophotometer (Agilent BioTek, BioTek Instruments Inc, USA). To verify RNA integrity, a 1% agarose gel electrophoresis was performed.

Reverse transcription was performed using the qScript cDNA synthesis kit, following the manufacturer’s protocols (Quantabio Reagent Technologies, QIAGEN Beverly Inc., USA) in a PCRmax Alpha Thermal cycler (cole-Parmer Ltd., UK). To synthesise the 20 µl final volume cDNA, we used a reaction mix containing qScript cDNA SuperMix, 200 ng total RNA and RNase/DNase-free water. The thermal cycling during cDNA synthesis was 25 °C for 5 min (priming), 42 °C for 30 min (reverse transcription) and 85 °C for 5 min (reverse transcriptase inactivation). The cDNA samples were diluted 10-fold and stored at −20 °C for Real-time PCR.

To measure mRNA expression, Quantitative real-time PCR (qPCR) was performed using HOT FIREPo EvaGreen qPCR mix Plus (Solis BioDyne, Teaduspargi, Estonia). Intron-spanning gene-specific primer pairs for quails were designed using Oligo7 software and obtained from Integrated DNA Technologies (BVBA-Leuven, Belgium) (see Supplementary Table S3 online for sequences of primers). We checked for target identity using Primer-Blast software of the National Centre for Biotechnology Information (NCBI) (http://www.ncbi.nlm.nih.gov). The qPCR was performed using the following thermal conditions: 95°C for 12 min (initial activation of the polymerase), 40 cycles of denaturation at 95°C for 15 s, annealing at 60 °C for 20 s and elongation at 72°C for 20 s. At the end of each run, the amplification specificity of each product was confirmed by melting curve analysis. Amplification and melting curve analysis (see supplementary Figure S1) and monitoring were performed using Agilent AreaMx Real-Time PCR System (Agilent Technologies, USA).

To identify a stable reference gene, we analysed commonly used genes in birds, namely beta-actin (*ACTB*), glyceraldehyde-3-phosphate dehydrogenase (*GAPDH*), and 18S ribosomal RNA (*RN18S*). We evaluated their stability and determined the most suitable reference gene, *ACTB*, by employing NormFinder, BestKeeper, and deltaCt algorithms^35^. The relative expression of mechanistic target of rapamycin (*mTOR*), ribosomal protein S6 kinase 1 (*RPS6K1*), autophagy-related 9A (*ATG9A*), growth hormone receptor (*GHR*), insulin-like growth factor 1 (*IGF1*) and its receptor (*IGF1R*) normalised to Beta-actin (*ACTB*) as reference gene was calculated using the efficiency corrected method^36^. The log of the expression ratio was used for statistical analyses of the relative mRNA expression, hereafter referred to as gene expression. To control for inter-plate variation, we repeated specific samples across plates for calibration. We used liver samples from male and female groups to analyse sex-specific expression. Samples from both sexes and all treatment groups were loaded on the same qPCR plate for each gene.

### Statistical analyses

All analyses were performed using R v. 4.2.2^38^. All images are processed using the ggplot function and saved at 300 DPI using the ggsave function, both from ggplot2 v.3.4.3 package ^39^. We fitted models to analyse our data depending on the data source and relationship of variables. Akaike’s information criterion corrected (AICc)^40^ was used to choose the best-supported models (Table S<u>4</u>, S<u>7</u>) and the final models utilised are described as follows. To analyse the effect of dietary treatment on body mass of males across restriction time, we employed linear mixed-effects models. Here, we considered dietary restriction with four levels, time points with three levels and their interaction as fixed factors, while individual bird identity as random intercept. We included individual bird identity as random factor to control for the effect of repeated measures^41^. We used the function ‘lmer’ from ‘lme4’ package^42^ to define both fixed and random effects and estimate model parameters. Additionally, we employed ‘lmerTest’ v. 3.1-3 package^43^ to compute p-values in ANOVA and model summary table (Table 1). We calculated mean body mass comparison of male birds among treatment groups within different time points (Table S<u>5</u>) and mean body mass comparison of male birds among time points (Table <u>S6</u>) within each treatment level using the function ‘emmeans’ with p < 0.05 significance level^44^.

**Table 1.** ANOVA output of the linear mixed-effect model indicating the effect dietary restriction on body mass across the two-week restriction period of male Japanese quails.

| Variables | Sum Sq | Mean Sq | NumDF | DenDF | F-value | p-value |
| --- | --- | --- | --- | --- | --- | --- |
| treatment | 629.9 | 209.96 | 3 | 28 | 2.96 | 0.049 |
| week | 3699.4 | 1849.71 | 2 | 56 | 26.08 | <.001 |
| treatment × week | 1781.7 | 296.96 | 6 | 56 | 4.19 | 0.002 |
| Random | K | logLik | AICc | LRT | df | p-value |
| 1\|birdID | 13 | -360.13 | 746.26 | 41.63 | 1 | <.001 |

Abbreviations: K, number of parameters; logLik, log-likelihood; AICc, Akaike’s information criterion corrected; LRT, Likelihood Ratio Test; df, degree of freedom.

We used linear models to analyse the effect of treatments on liver gene expression in male Japanese quails. In this model, the expression of each gene was treated as a response variable, while the dietary treatments served as the explanatory variable. Statistical significance was assessed through one-way ANOVA (Table 3), and the means of the treatment group were computed using the function ‘emmeans’ with p < 0.05 significance level (Table <u>S9</u>). We performed a principal component analysis (PCA), to reduce the dimensionality of the gene expression data, to transform the original correlated gene expression variables into a set of linearly uncorrelated principal components. This was applied using ‘prcomp’ function from the ‘stats’ package^38^. PCA was used to avoid multicollinearity occurring between the predictor variables (gene expressions). We used ‘ggbiplot’ package^45^ to visualise the pattern of the variables (gene expression) against the treatments groups. Furthermore, we used Kaiser’s rule to select which PCs to retain for subsequent regression analysis^46^. Finally, the first two components, PC1 and PC2, met the Kaiser’s rule and were used in linear regression against body mass (Table <u>S11</u>).

To compute differences between males and females and compare sex-specific responses to dietary restriction, we included previously reported data from a parallel experiment conducted in females, following the same protocols and conditions^32^. Using the male and female data, we employed linear mixed-effect models to analyse the sex-specific effects of dietary restriction on body mass, where sex, dietary treatment, time points and their interactions considered as fixed factors and individual bird identity as random factor. We used three-way ANOVA to analyse statistical significance (Table 2) and ‘emmeans’ to compute mean body mass of males and females at different dietary restriction levels across the time points (Table <u>S8</u>). We utilised linear models to analyse the sex-specific effects of dietary restriction on liver gene expression. We assessed the significance of the effects of sex, dietary treatment, and their interaction using a two-way ANOVA (Table 4) and means of males and females within treatment groups were computed using ‘emmeans’ with p < 0.05 significance level (Table S<u>10</u>). The experimental block did not significantly contribute to any of the models; therefore, we removed it from all the final models.

**Table 2.** Output of three-way ANOVA of the linear mixed-effect model, (body mass ∼ treatment × sex × week), indicating the effects of dietary restriction on body mass across the two-week restriction period.

| Variables | Sum Sq | Mean Sq | NumDF | DenDF | F-value | p-value |
| --- | --- | --- | --- | --- | --- | --- |
| treatment | 3766.2 | 1255.4 | 3 | 56 | 14.70 | <.001 |
| sex | 4987.7 | 4987.7 | 1 | 56 | 58.41 | <.001 |
| week | 16065.0 | 8032.5 | 2 | 112 | 94.06 | <.001 |
| treatment × sex | 959.0 | 319.7 | 3 | 56 | 3.74 | 0.016 |
| treatment × week | 8456.5 | 1409.4 | 6 | 112 | 16.50 | <.001 |
| sex × week | 1687.6 | 843.8 | 1 | 112 | 9.88 | <.001 |
| treatment × sex × week | 1542.0 | 257.0 | 6 | 112 | 3.01 | 0.009 |
| Random | K | logLik | AICc | LRT | df | p-value |
| 1\|birdID | 25 | -737.67 | 1525.3 | 85.39 | 1 | <.001 |

Abbreviations: K, number of parameters; logLik, log-likelihood; AICc, Akaike’s information criterion corrected; LRT, Likelihood Ratio Test; df, degree of freedom.

### Ethical approval and consent to participate

The experiment was performed following the EU Directive “Legislation for the protection of animals used for scientific purposes” and after approval by the Ethical Committee for Animal Use of the University of Debrecen, Hungary (Protocol No. 5/2021/DEMAB). We confirm that all the methods were carried out in compliance with relevant institutional guidelines and regulations. Our research findings are presented following the ARRIVE guidelines.

## Results

### Only severe dietary restriction reduces male body mass

The dietary restriction (DR) and its interaction with restriction period (weeks) had a significant effect on body mass in male quails (Table 1). All male quails from restricted groups showed a reduced trend in body mass compared to the quails from ADL group (Fig. 1). However, only quails from DR40 proved a statistically significant reduction in the first and second weeks (week 1: *p* = 0.02; week 2: *p* < 0.001; Table <u>S5</u>). When compared to their initial body mass, all male quails from restricted groups showed significantly reduced body mass on both week 1 and week 2, whereas only male quails from DR40 groups showed further mass reduction from week 1 to week 2 (*p* = 0.050; Table <u>S6</u>).

**Figure 1.**
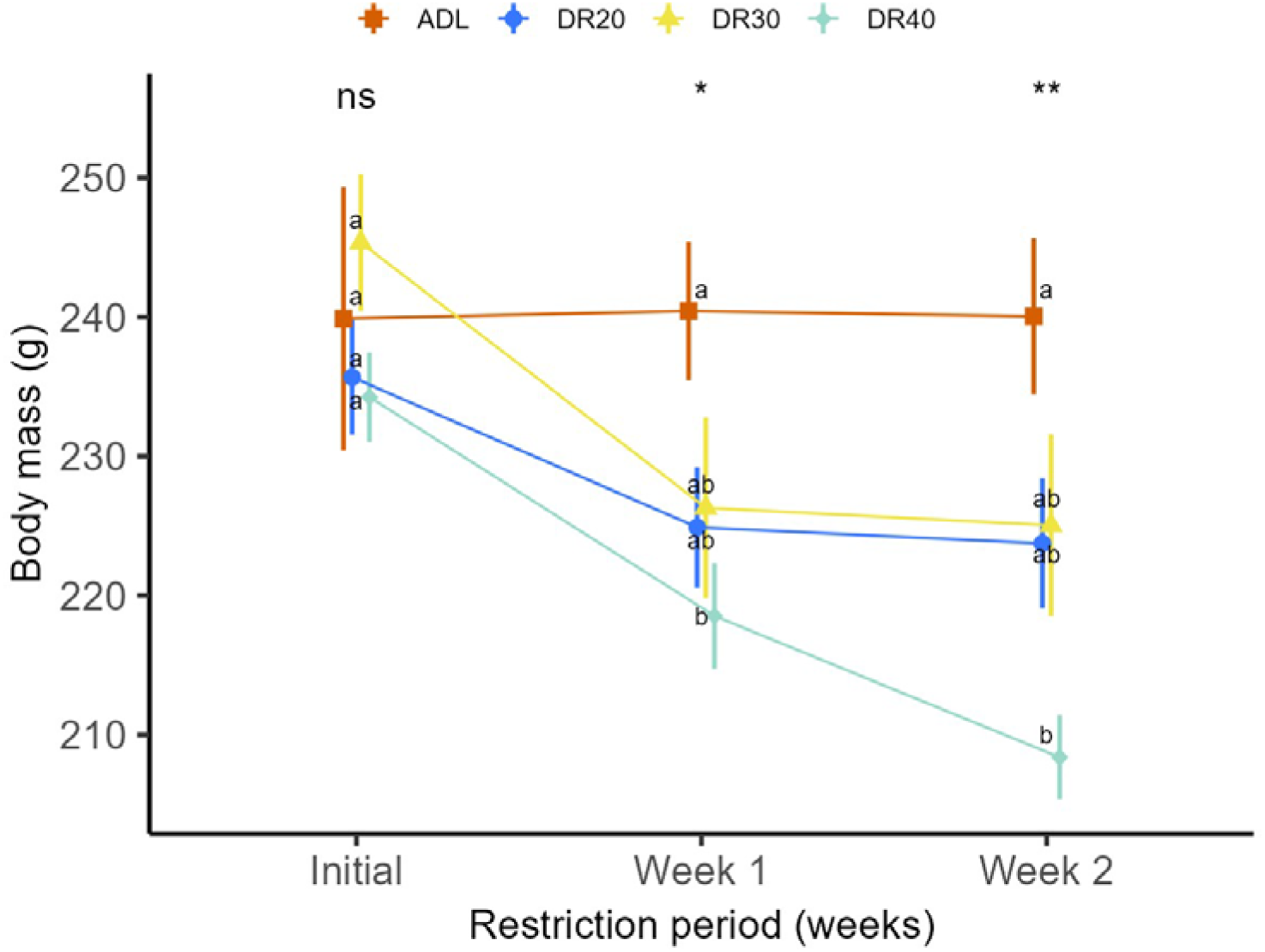
The effect of different dietary restriction levels on body mass of male Japanese quails across the two-week restriction period. Data are represented by the meanLJ±LJSEM from 8 birds per group and were analysed using two-way ANOVA from linear mixed effect model. Means followed by common letters within time points are not significantly different at p < 0.05. Abbreviations: ADL, *ad libitum*; DR20, 20% restriction; DR30, 30% restriction; DR40, 40% restriction. Initial, day 0; week 1, day 7; week 2, day 14. ‘ns’, not significant at p < 0.05; ‘*’ significantly different at p < 0.05; ‘**’ significantly different at p < 0.01

### Dietary restriction exerts sex-specific effect on body mass

Over time, males and females exhibited distinct responses to dietary restriction, as indicated by a significant interaction between sex, treatment, and the restriction period (Table 2). Males showed significantly lower initial body mass than females in all treatment groups, and this difference persisted throughout the experiment in the ADL fed and moderately restricted (DR20) groups (Fig. 2, Table <u>S8</u>). However, the difference in body mass between males and females disappeared by the second week in the DR30 group and by the first week in the DR40 group (Fig. 2, Table S<u>8</u>).

**Figure 2.**
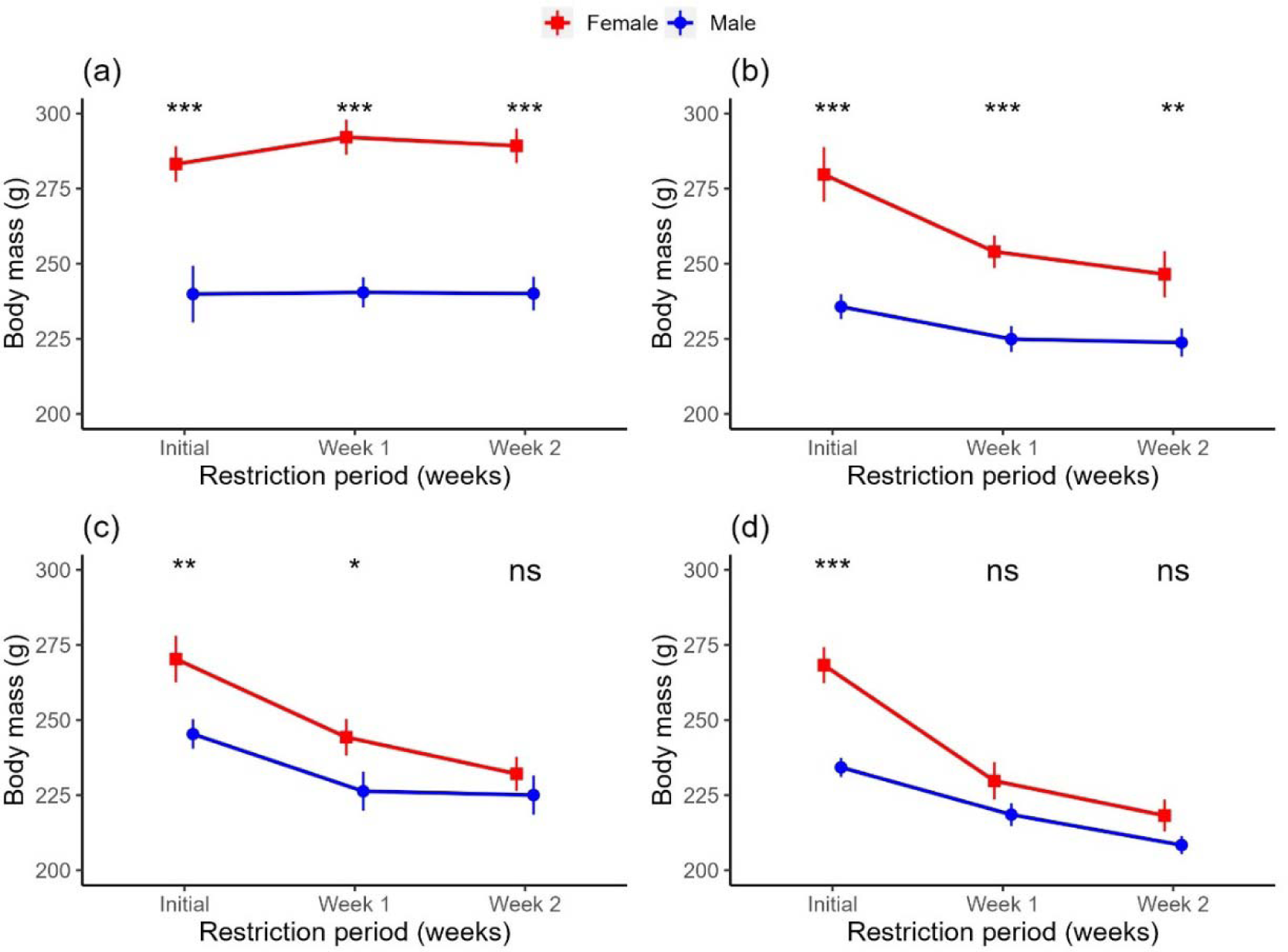
Comparing body mass of female and male Japanese quails in different dietary restriction levels across restriction period. (a) ADL, *ad libitum* group, (b) DR20, 20% restriction (c) DR30, 30% restriction, (d) DR40, 40% restriction . Dots and vertical bars represent the meanLJ±LJSEM from 8 birds per group, and data were analysed using ANOVA of linear mixed-effect model. Female data is obtained from our previous study^32^. Abbreviations: ‘ns’, not significant at p < 0.05; ‘*’ significantly different at p < 0.05; ‘**’ significantly different at p < 0.01; *** significantly different at p < 0.001.

### Dietary restriction affects mTOR signalling genes in the liver of males

The linear model indicated that DR significantly affected the expression of major genes responsible for liver mTOR signalling pathway in males (Table 3). Compared to the ADL group, all restricted groups showed significantly lower expression of hepatic *mTOR* and *IGF1* genes. However, there was no significant variation among the restricted groups on both genes (Fig. 3a,d; Table <u>S9</u>). The expression of *IGF1R* also decreased with increasing severity of restriction, with the DR40 group showing a significantly lower value (*p* = 0.009) compared to the ADL group and a marginally lower value (*p* = 0.074) compared to the DR30 group (Fig. 3e, Table <u>S9</u>). Additionally, DR increased the expression of *RPS6K1* and *ATG9A* genes (Fig. 3b,c, Table <u>S9</u>). In both genes, all the restricted groups showed significantly higher expression than the ADL group, while there was no significant difference among the restricted groups. While ANOVA showed significant treatment effect on *GHR* expression (Table 3), the pairwise comparisons showed no significant differences among treatment groups (Fig. 3f, Table <u>S9</u>).

**Table 3.** ANOVA output of a linear model, lm(gene ∼ treatment), showing the effect of dietary restriction treatment on expression of key mTOR pathway genes in male Japanese quails.

| Explanatory variable | Gene (response) | Sum Sq | Mean Sq | df | F-value | p-value |
| --- | --- | --- | --- | --- | --- | --- |
| Treatment | *mTOR* | 4.77 | 1.59 | 3 | 6.46 | 0.002 |
|  | *RPS6K1* | 30.39 | 10.13 | 3 | 14.01 | <.001 |
|  | *ATG9A* | 5.29 | 1.76 | 3 | 7.22 | <.001 |
|  | *IGF1* | 30.44 | 10.15 | 3 | 15.11 | <.001 |
|  | *IGFR* | 8.56 | 2.85 | 3 | 4.28 | 0.013 |
|  | *GHR* | 6.75 | 2.25 | 3 | 3.61 | 0.025 |

Abbreviations: *mTOR*, mechanistic target of rapamycin; *RPS6K1*, ribosomal protein S6 kinase 1; *ATG9A*, autophagy-related 9A; *IGF1*, insulin-like growth factor 1; *IGF1R*, insulin-like growth factor 1 receptor; *GHR*, growth hormone receptor;

**Figure 3.**
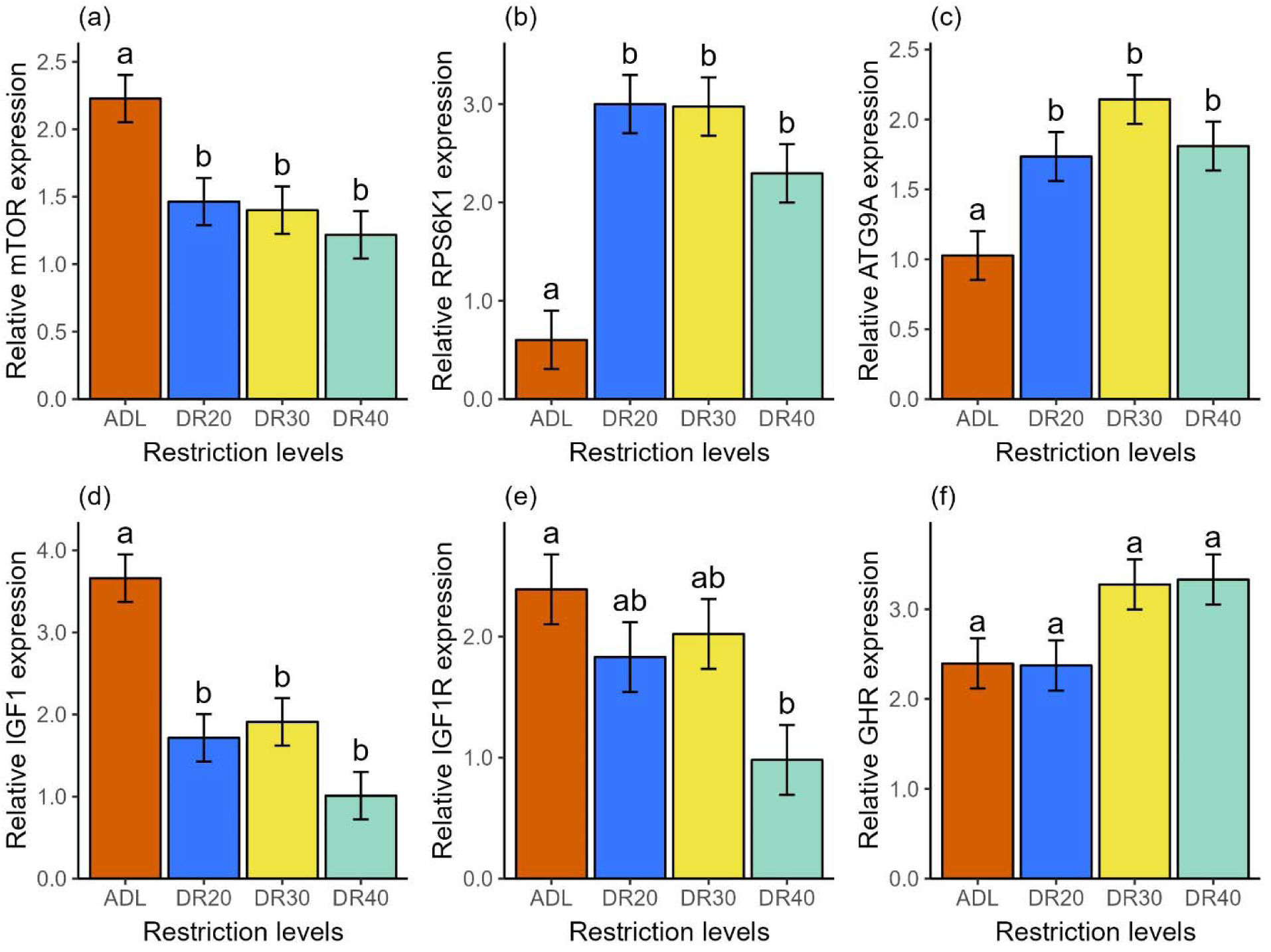
Effects of dietary restriction on expression of genes mediating nutrient availability in male Japanese quails. (a) *mTOR,* mechanistic target of rapamycin, (b) *RPS6K1,* ribosomal protein S6 kinase 1, (c) *ATG9A,* autophagy-related 9A, (d) *IGF1,* insulin-like growth factor 1, (e) *IGF1R,* insulin-like growth factor 1 receptor, (f) *GHR,* growth hormone receptor. Relative mRNA expression is analysed in log of fold change. Data are represented by the meanLJ±LJSEM from 8 birds per group. The Tukey test was used as a *post hoc* test at p < 0.05 significance level. Means followed by a common letter are not significantly different at p < 0.05. Abbreviations: ADL, *ad libitum*; DR20, 20% restriction; DR30, 30% restriction; DR40, 40% restriction.

### Liver mTOR signalling genes showed sex-specific expression intensity

In order to explore how genes respond differently based on sex, we combined gene expression data from both males and females. As a result, the expression levels of the liver *mTOR* pathway genes showed significant variation between male and female groups in response to varying levels of DR over a period of two weeks (Table 4). Females showed higher *mTOR* expression than the males in the ADL (*p* =0.004) and DR20 (*p* = 0.018) groups, while no significant difference was observed between the sexes in the severely restricted groups (Fig. 4a, Table <u>S10</u>). Females exhibited higher *RPS6K1* expression in the ADL fed group (*p* = 0.080), while lower in the DR20 (*p* = 0.003) and DR30 (*p* = 0.087) groups (Fig. 4b, Table <u>S10</u>). *ATG9A* showed lower expression in females compared to males, with significant differences observed at DR20 (*p* < 0.005) and DR30 (*p* < 0.001) groups (Fig. 4c, Table <u>S10</u>). Females showed higher *IGF1* expression, with significant value in the ADL (p = 0.003), DR20 (p = 0.024) and DR40 (*p* <.001) groups (Fig. 4d, Table <u>S10</u>). Furthermore, females exhibited higher *IGF1R* expression than the males at the severely restricted groups (*p* = 0.048; Fig. 4e, Table <u>S10</u>), while *GHR* showed no significant variation between the sexes (Fig. 4F, Table <u>S10</u>).

**Table 4.** Output of two-way ANOVA from a linear model, *lm(gene ∼ treatment × sex)*, indicating the effect of dietary restriction on expression of liver mTOR pathway genes across the two-week restriction period.

| Response variables | Explanatory Variables | Sum Sq | Mean Sq | df | F-value | p-value |
| --- | --- | --- | --- | --- | --- | --- |
| *mTOR* | treatment | 22.92 | 7.64 | 3 | 20.83 | <.001 |
|  | sex | 1.84 | 1.84 | 1 | 5.02 | 0.029 |
| | treatment × sex | 3.84 | 1.28 | 3 | 3.49 | 0.021 |
| *RPS6K1* | treatment | 25.62 | 10.05 | 3 | 18.94 | <.001 |
|  | sex | 0.75 | 0.75 | 1 | 1.59 | 0.212 |
| | treatment × sex | 6.48 | 2.16 | 3 | 479 | 0.005 |
| *ATG9A* | treatment | 6.06 | 2.02 | 3 | 8.17 | <.001 |
|  | sex | 5.34 | 5.34 | 1 | 21.59 | <.001 |
| | treatment × sex | 2.18 | 0.73 | 3 | 2.93 | 0.041 |
| *IGF1* | treatment | 57.22 | 19.07 | 3 | 19.80 | <.001 |
|  | sex | 32.79 | 32.79 | 1 | 34.05 | <.001 |
| | treatment × sex | 7.01 | 2.34 | 3 | 2.42 | 0.075 |
| *IGF1R* | treatment | 6.71 | 2.24 | 1 | 4.13 | 0.010 |
|  | sex | 0.01 | 0.01 | 3 | 0.02 | 0.895 |
| | treatment × week | 3.17 | 1.06 | 1 | 1.95 | 0.132 |
| *GHR* | treatment | 25.56 | 8.52 | 3 | 11.09 | <.001 |
|  | sex | 0.71 | 0.71 | 1 | 0.93 | 0.340 |
| | treatment × sex | 2.12 | 0.77 | 3 | 0.92 | 0.437 |

Abbreviations: *mTOR*, mechanistic target of rapamycin; *RPS6K1*, ribosomal protein S6 kinase 1; *ATG9A*, autophagy-related 9A; *IGF1*, insulin-like growth factor 1; *IGF1R*, insulin-like growth factor 1 receptor; *GHR*, growth hormone receptor.

**Figure 4.**
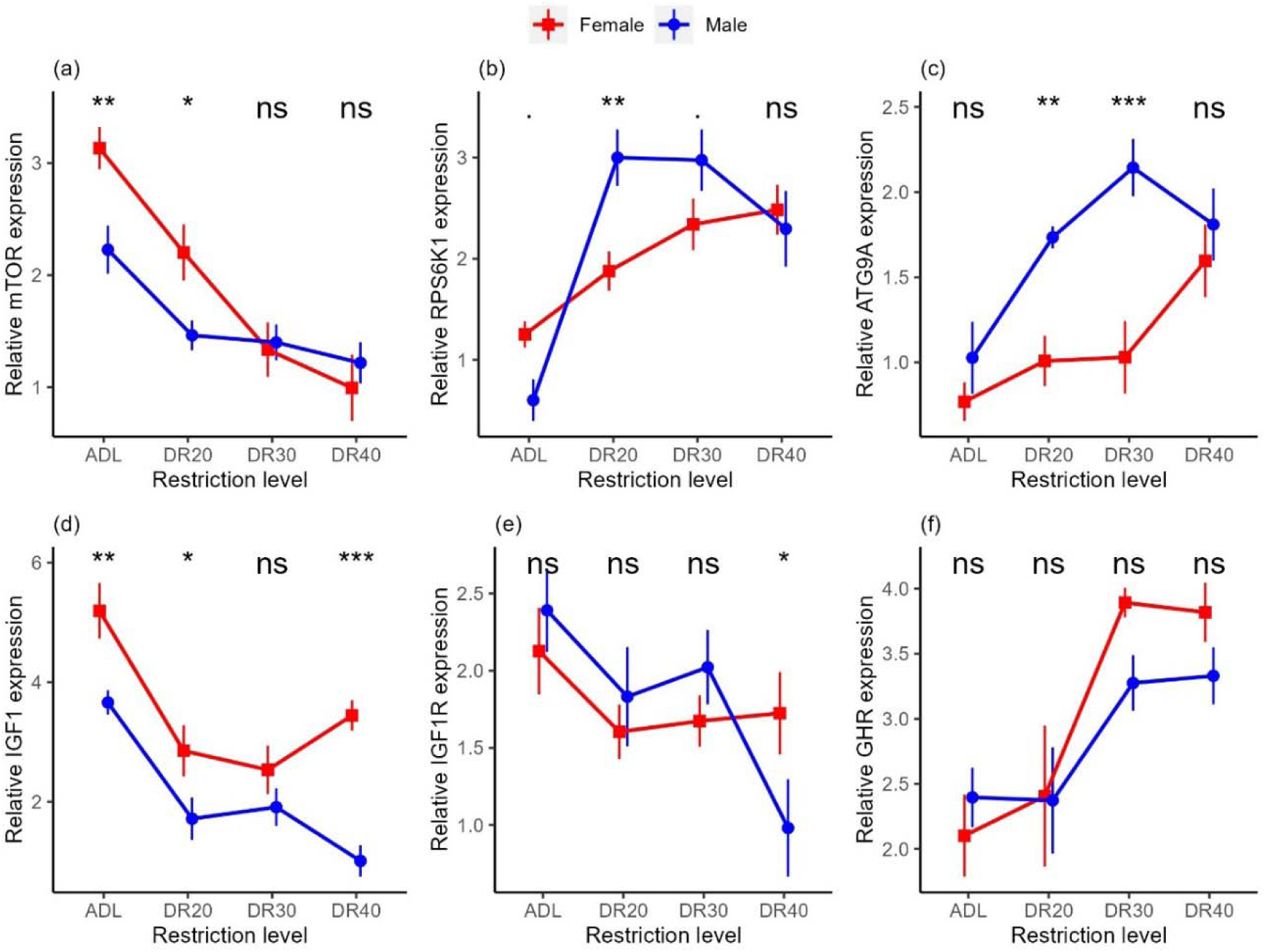
Sex-specific effects of dietary restriction in Japanese quails. (a) *mTOR,* mechanistic target of rapamycin; (b) *RPS6K1,* ribosomal protein S6 kinase 1; (c) *ATG9A,* autophagy-related 9A, (d) *IGF1,* insulin-like growth factor 1; (e) *IGF1R,* insulin-like growth factor 1 receptor; (f) *GHR,* growth hormone receptor. Relative mRNA expression is analysed in log fold change. Dots and vertical bars represent the meanLJ±LJSEM from 8 birds per group and data were analysed using ANOVA of linear model. Abbreviations: ADL, *ad libitum*; DR20, 20% restriction; DR30, 30% restriction; DR40, 40% restriction. ‘ns’, not significant at p < 0.05; ‘.’ marginally insignificant. ‘*’ significantly different at p < 0.05; ‘**’ significantly different at p < 0.01; ‘***’ significantly different at p < 0.001.

### Principal components are associated with variation in body mass

To disentangle the complex interplay of liver gene expression of male quails, we employed principal component analysis (PCA). The PCA indicated that the first two principal components have eigenvalues greater than 1 (Table <u>S11</u>) and thus were retained for further regression analysis. These two components explained 69.0% (PC1 = 42.1% and PC2 = 26.9%) of the total variance. The analysis revealed that expression of *mTOR*, *IGF1*, *RPS6K1* and *ATG9A* contribute significantly to PC1, influencing variation in different directions, while expression of *IGF1R* and *GHR* predominantly shaped PC2 (Table <u>S12</u>). The elliptical biplot indicated a clustering of liver *mTOR* and *IGF1* genes expression around the control treatment, while *RPS6K1* and *ATG9A* genes expression clustered around the groups received restricted treatments, aligning with their positive and negative influences on PC1, respectively (Fig. 5). Finally, we showed that PC1 showed a positive correlation with body mass, indicating a potential link between gene expressions resembling the control treatment and the body mass of quails after two weeks of DR. Conversely, PC2 displayed a negative association with body mass, suggesting a contrasting impact associated with gene expressed on the restricted groups (Table 5).

**Figure 5.**
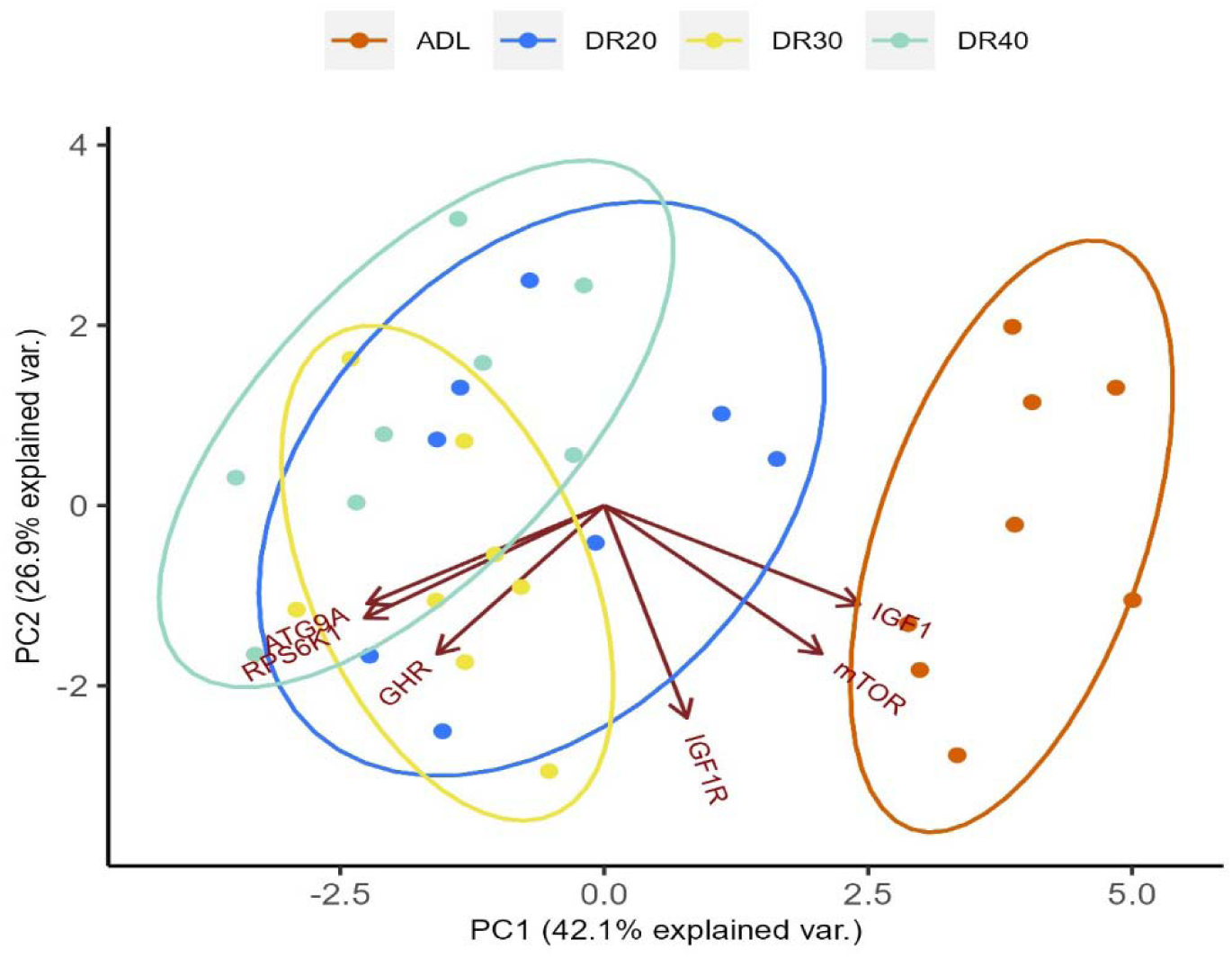
A biplot of PCA for the liver gene expression and body mass in Japanese quails treated with different dietary restriction levels for two weeks. Clustering is based on dietary restriction levels and a dimensional indication of genes in line with the restriction levels. The ellipsoids are defined by the treatment groups. Abbreviations: ADL, *ad libitum*; DR20, 20% restriction; DR30, 30% restriction; DR40, 40% restriction; *mTOR,* mechanistic target of rapamycin; *RPS6K1,* ribosomal protein S6 kinase 1; *ATG9A,* autophagy-related 9A; *IGF1,* insulin-like growth factor 1; *IGF1R,* insulin-like growth factor 1 receptor; *GHR,* growth hormone receptor.

**Table 5.** Summary of multiple linear regression model to estimate the change in body mass of male Japanese quails with principal components.

| Predictors | estimate | SEM | t-value | p-value |
| --- | --- | --- | --- | --- |
| Intercept | 224.40 | 3.23 | 69.54 | <.001 |
| PC1 | 4.89 | 2.04 | 2.39 | 0.025 |
| PC2 | -5.45 | 2.41 | -2.26 | 0.034 |
| df | 24 |  |  |  |
| R<sup>2</sup> | 0.33 |  |  |  |
| RSE | 16.13 |  |  |  |
| F-stat | 5.42 |  |  |  |
| Cumulative P-value | 0.012 |  |  |  |

## Discussion

Nutritional availability plays a crucial role in shaping an individual’s phenotype in response to environmental conditions. The intricate interplay between an organism’s genetic makeup and the availability of essential nutrients profoundly influences its overall fitness. This dynamic relationship has the power to modulate gene expression, thereby shaping phenotypic outcomes in various environmental contexts ^47,48^. In natural conditions, organisms often face challenges such as food shortages, prompting them to make necessary molecular and physiological adjustments to cope with these adversities. Imbalances in nutrient intake can lead to altered gene expression patterns across various tissues, particularly the liver, thereby contributing to the plasticity of fitness outcomes^49–51^. In response to environmental cues, the expression levels of genes contribute to either the upregulation or downregulation of their respective proteins/peptides. Consequently, these proteins/peptides mediate the information received from variations in gene expression to influence fitness plasticity ^52,53^. In the face of changing dietary conditions, liver signalling pathways undergo substantial alterations in gene expression levels, representing an adaption mechanism^51,54^. These transcriptional changes implicate a broad spectrum of biological processes including nutrient metabolism, protein synthesis and detoxification, thereby governing phenotypic plasticity^51,55^. Notably, nearly every gene involved in the mTOR signalling pathway demonstrates differential expression in the liver, exerting a profound influence on the overall functionality of the organisms. This ensures efficient utilisation of available nutrients^33,56^ Organisms could show sex-specific responses to nutritional variability because of differences in reproductive investment^1,30^. Previously, in female Japanese quails we found that body mass and expression of genes mediating the mTOR signalling pathway are affected differently at different dietary restriction (DR) levels^32^. In the current study, we used an identical protocol to investigate the effect of different levels of DR on male Japanese quails, expecting a different response between males and females to nutrient availability. We found a significant effect of DR on hepatic gene expression, largely consistent with the picture in females, albeit different in magnitude for certain genes.

Despite a similar trend of change with females in all restricted groups, only the severely restricted (DR40) group showed significantly reduced body mass compared to the control group on both week 1 and week 2 (Fig. 1). The resource allocation strategies of male and female birds are shaped by their different reproductive roles and selective pressures. Female birds typically invest more resources in reproduction than males. In contrast, males usually focus more on attracting mates and competing with other males for access to females^57,58^. Therefore, the observed higher mass loss in females due to dietary restrictions may be due to their substantial investment in egg production, since the birds were laying eggs during our study^32^. The variation could be regulated by the mechanism of expression of genes governing nutrient-sensing pathway.

The *IGF1* and its receptors (*IGF1R*) are the crucial genes of interest in a nutrient-sensing pathway^59–61^. IGF-1 and its receptor, IGF1R, play pivotal roles in growth and development across a wide spectrum of organisms. IGF-1, mainly secreted in the liver, acts as a critical mediator of cell growth, differentiation, and survival. It exerts its effects by binding to its specific receptor, that triggers a cascade of signalling, mainly mTOR pathway, a pathway responsible for cellular proliferation ^62,63^. The IGF-1/IGF1R axis also plays a pivotal role in sexual dimorphism, influencing the distinct physiological and morphological differences between males and females upon interaction with sex steroids^64–66^. In both sexes, expression of *IGF1* and *IGF1R* genes was reduced in all restriction levels compared to the ADL group (Fig. 4d,e). The trend in *IGF1* gene expression across the DR gradients in males mirrors our findings in females, albeit with lower expression levels in male groups. In the case of *IGF1R* gene expression, the pattern of change is similar in both sexes up to DR30 level, although males showed higher reduction at the severe restriction level (DR40) (Fig. 4e). Although little is known about the sex differences on the impact of DR on *IGF1* gene expression, previous studies examining circulating IGF-1 have suggested that both the level and the influence of IGF-1 exhibit sex-based difference in mammals^65^ and in birds^64,67–69^. For instance, a study in chickens suggested a strong correlation between the plasma levels of IGF-1 and expression of the *IGF1* gene in the liver^70–73^. Therefore, the evidence indicating sex-specific level of plasma IGF-1 could align with the hepatic gene expression patterns we observed. Reduced *IGF1* gene expression due to DR also suggests a corresponding effect on circulating IGF-1 levels^71,72^. Therefore, the current study suggests sex-specific *IGF1* gene expression but not treatment-specific differences between females and males.

We also tested the effect of DR on growth hormone receptor (*GHR*), a receptor protein that binds to the growth hormone to initiate *IGF1* expression (Fig. 3,4, Table 3<u>,4</u>). Contrary to findings in mammals and fish, where nutritional deficit reduces *GHR* expression and produces growth hormone resistance to limit *IGF1* expression^74–76^ ^77,78^, our results showed that *GHR* gene expression was significantly upregulated in the severely restricted groups in females while remaining marginal in males. The higher expression in females compared to males in the DR40 group coincided with what we observed in *IGF1* expression, where males downregulated *IGF1* expression more than did females. Feed restriction also increased *GHR* expression in chicken^79^. Therefore, the nutritional regulation of the hypothalamo-pituitary-somatotropic axis may be different in birds, which requires further research.

Another gene of interest within the nutrient-sensing pathway is the *mTOR*, which plays a role in nuclear transcription and lysosomal translation. The impact of DR on the mTOR signalling pathway, a crucial molecular marker in nutrient sensing, was significant. This effect is achieved by modulating the expression of specific genes based on their function^80,81^. The present study in male quails revealed that in comparison to the ADL group, *mTOR* gene expression was decreased similarly across all restriction levels (Fig. 3a). This pattern is different from that of in females, where the reduction of *mTOR* expression intensified with increased levels of restriction. Furthermore, while females exhibit significantly higher *mTOR* expression in the *ad libitum*-fed and moderately restricted (DR20) groups than males, the difference between the sexes disappeared in the severely restricted (DR30 and DR40) groups (Fig. 4a). This indicates that similar to body mass (Fig. 2), the effect of DR was stronger on female birds than on males, which may be due to physiological, morphological, and hormonal differences and reproduction strategies between the sexes^21,82,83^. The females’ need for more food than males to maintain the larger body mass and egg production may force their *mTOR* expression to respond strongly to restriction. This is evidence to suggest that the *mTOR* gene expression mediates the effect of DR on body mass. The principal components analysis also provides evidence that *mTOR* is one of the major contributors to PC1, which positively explains body mass in both males (Fig. 5, Table 5) and females^32^. Notably, the association appears to be stronger in females.

Another gene downstream of *mTOR* is *ATG9A*, a gene initiating the formation of autophagosomes for the degradation of cellular contents in response to nutritional deficiency^83–85^. Our finding demonstrated that the expression of the *ATG9A* gene was upregulated in all the restricted groups compared to the ADL-fed group in both females and males (Fig. 4c), in stark contrast to the effect observed on the *mTOR* and *IGF1* genes. The *ATG9A* expression level was higher in males in the DR20 and DR30 groups. The effect of DR is consistent with previous studies from other organisms^86,87^. However, the pattern of changes across restriction levels was different: males showed a pronounced increase at all restriction levels, while females showed a significant increase only at the severely restricted level, resulting in significant sex differences in the DR20 and DR30 groups (Fig. 4c). At the expense of anabolic progressions and stress, DR has a critical role in maintaining pathways required to retain cellular function. In conditions of scarce resources, autophagy serves as a cytoprotective mechanism through recycling of damaged organelles and malformed proteins, a process in which cells break down their components to provide energy and nutrients^88,89^. DR upregulates autophagy, one way, through inhibition of mTOR activity that facilitates the nuclear localisation of Transcription Factor EB (TFEB), the ATG transcription factors in the nucleus^90,91^. Accordingly, the lower *mTOR* expression in both males and females may have a significant contribution to the upregulation of *ATG9A* expression (Fig. <u>S2</u>). However, the pronounced increase in *ATG9A* expression, specifically in males, may contribute to the rapid recycling of cytoplasmic waste and supply it as energy and amino acids for other cellular activities. The process could potentially contribute to keep the pace of *mTOR* expression through a positive feedback mechanism, as we observed sustained expression levels across all restriction groups and relatively lower body mass loss in males. In females, there was a moderate increase in *ATG9A* expression and a rapid reduction in *mTOR*.

The other gene of interest we studied is the *RPS6K1*, a gene situated downstream of mTOR. Our study showed that DR upregulates the expression of *RPS6K1* in both males and females (Fig. 4b). The result contradicted our assumption that DR would have a downregulating role. The pattern of increase was more pronounced in males in the DR20 and DR30 groups. Since RPS6K1 plays a critical role in ribosomal translation^92,93^, its gene expression would contribute to maintaining the level of the respective kinase for phosphorylation. This, in turn, would help reduce the loss of body mass that we observed in females but less in males.

In conclusion, our study revealed that dietary restriction affects body mass and the expression of critical genes governing the mTOR pathway in male Japanese quails. These results corroborate the general gene expression patterns seen in females^32^. However, for the first time in birds, we provide evidence that the fine-scale regulation of the mTOR pathway is sex-specific, as seen in the differential expression of most of the genes studied. Female birds exhibited higher body mass and more intensive mass loss than males and demonstrated intensified reduction in *mTOR* gene expression with increasing restriction levels. These findings align with females’ larger body size and reproductive investment. In contrast, males exhibited a more pronounced upregulation in *ATG9A* gene expression, potentially aiding their ability to avoid severe body mass loss. This suggests that males may have a higher capacity for cellular waste recycling and energy utilisation under dietary restriction conditions. The anabolic genes of *RPS6K1* and *GHR* showed sex-specifically intensified expression in the restricted groups, contrary to our assumption. These sex-specific responses shed light on the intricate interplay between nutrient availability, gene expression, and body size, highlighting the importance of considering sexual dimorphism in studies of dietary restriction, and animal physiology in general.

## Acknowledgements

We thank three anonymous referees who provided constructive comments on an earlier version of this manuscript. We would like to thank Angyal Eszter and Fadella Nur Almira for their help during the experiment.

## Data availability

All data generated or analysed during this study are included in this published article [and its supplementary information files].

Upon acceptance of the manuscript, the datasets will be uploaded into a public repository.

## Author contributions

G.K.R.: conceptualisation, data curation, formal analysis, investigation, visualisation, methodology, validation, writing—original draft, writing—review and editing; S.F.N.: conceptualisation, methodology, writing—review and editing; B.C.: methodology and writing— review and editing; R.K.: methodology and writing—review and editing; C.S.: methodology, resource and writing—review and editing; J.K.L.: methodology and writing—review and editing; L.C.: conceptualisation, supervision, funding acquisition, project administration, resources, validation and writing—review and editing; A.Z.L.: conceptualisation, methodology, supervision, funding acquisition, project administration, resources, validation and writing— review and editing.

All authors gave final approval for publication and agreed to be held accountable for the work performed therein.

## Funding

The study was funded by the National Development, Research and Innovation Office (OTKA K139021).

## Competing interests

The authors declare that there are no competing interests.

## Additional information

Supplementary Information is submitted in a separate file.

## Notes

### Competing Interest Statement

The authors have declared no competing interest.

## References

1 Gegenhuber, B., Wu, M., Bronstein, R. & Tollkuhn, J. Gene regulation by gonadal hormone receptors underlies brain sex differences. Nature 606, 153–159. 10.1038/s41586-022-04686-1 (2022).

2 Arlettaz, R., Christe, P. & Schaub, M. Food availability as a major driver in the evolution of life_history strategies of sibling species. Ecology and Evolution 7, 4163–4172. 10.1002/ece3.2909 (2017).

3 Johnson, S. C. “Nutrient sensing, signaling and ageing: The role of IGF-1 and mTOR in ageing and age-related disease” in Biochemistry and Cell Biology of Ageing: Part I Biomedical Science Vol. 90 Subcellular Biochemistry 49–97 (Springer Singapore, 2018).

4 Mc Auley, M. T. Dietary restriction and ageing: recent evolutionary perspectives. Mechanisms of Ageing and Development 208, 111741. 10.1016/j.mad.2022.111741 (2022).

5 Rollins, J. A., Shaffer, D., Snow, S. S., Kapahi, P. & Rogers, A. N. Dietary restriction induces posttranscriptional regulation of longevity genes. Life Science Alliance 2, e201800281. 10.26508/lsa.201800281 (2019).

6 Nourmohammad, A. et al. Adaptive evolution of gene expression in Drosophila. Cell reports 20, 1385–1395. 10.1016/j.celrep.2017.07.033 (2017).

7 Efeyan, A., Comb, W. C. & Sabatini, D. M. Nutrient-sensing mechanisms and pathways. Nature 517, 302–310. 10.1038/nature14190 (2015).

8 Laplante, M. & Sabatini, D. M. mTOR signaling in growth control and disease. Cell 149, 274–293. 10.1016/j.cell.2012.03.017 (2012).

9 Regan, J. C., Froy, H., Walling, C. A., Moatt, J. P. & Nussey, D. H. Dietary restriction and insulin_like signalling pathways as adaptive plasticity: A synthesis and re_evaluation. Functional Ecology 34, 107–128. 10.1111/1365-2435.13418 (2020).

10 Kennedy, B. K. & Lamming, D. W. The mechanistic target of rapamycin: the grand conducTOR of metabolism and aging. Cell Metabolism 23, 990–1003. 10.1016/j.cmet.2016.05.009 (2016).

11 Sandri, M. et al. Signalling pathways regulating muscle mass in ageing skeletal muscle. The role of the IGF1-Akt-mTOR-FoxO pathway. Biogerontology 14, 303–323. 10.1007/s10522-013-9432-9 (2013).

12 Papadopoli, D. et al. mTOR as a central regulator of lifespan and aging. F1000Research 8, 998. 10.12688/f1000research.17196.1 (2019).

13 Gollwitzer, P., Grützmacher, N., Wilhelm, S., Kümmel, D. & Demetriades, C. A Rag GTPase dimer code defines the regulation of mTORC1 by amino acids. Nature Cell Biology 24, 1394–1406. 10.1038/s41556-022-00976-y (2022).

14 Hara, K. et al. Amino acid sufficiency and mTOR regulate p70 S6 kinase and eIF-4E BP1 through a common effector mechanism. Journal of Biological Chemistry 273, 14484–14494. 10.1074/jbc.273.23.14484 (1998).

15 Laplante, M. & Sabatini, D. M. Regulation of mTORC1 and its impact on gene expression at a glance. Journal of Cell Science 126, 1713–1719. 10.1242/jcs.125773 (2013).

16 Hansen, M. et al. Lifespan extension by conditions that inhibit translation in Caenorhabditis elegans. Aging cell 6, 95–110. 10.1111/j.1474-9726.2006.00267.x (2007).

17 Kabil, H., Kabil, O., Banerjee, R., Harshman, L. G. & Pletcher, S. D. Increased transsulfuration mediates longevity and dietary restriction in Drosophila. PNAS 108, 16831–16836. 10.1073/pnas.110200810 (2011).

18 Chen, C.-N., Liao, Y.-H., Tsai, S.-C. & Thompson, L. V. Age-dependent effects of caloric restriction on mTOR and ubiquitin-proteasome pathways in skeletal muscles. GeroScience 41, 871–880. 10.1007/s11357-019-00109-8 (2019).

19 Holmes, D. J. & Ottinger, M. A. Birds as long-lived animal models for the study of aging. Experimental Gerontology 38, 1365–1375. 10.1016/j.exger.2003.10.018 (2003).

20 Satoh, T. Bird evolution by insulin resistance. Trends in Endocrinology & Metabolism 32, 803–813. 10.1016/j.tem.2021.07.007 (2021).

21 Bennett-Keki, S., Fowler, E. K., Folkes, L., Moxon, S. & Chapman, T. Sex-biased gene expression in nutrient-sensing pathways. Proceedings of the Royal Society B 290, 20222086. 10.1098/rspb.2022.2086 (2023).

22 McDonald, J. M. C., Nabili, P., Thorsen, L., Jeon, S. & Shingleton, A. W. Sex-specific plasticity and the nutritional geometry of insulin-signaling gene expression in Drosophila melanogaster. EvoDevo 12, 6. 10.1186/s13227-021-00175-0 (2021).

23 Khodursky, S., et al. Sex differences in interindividual gene expression variability across human tissues. PNAS Nexus 1, pgac243. 10.1093/pnasnexus/pgac243 (2022).

24 Bazhan, N. et al. Sex differences in liver, adipose tissue, and muscle transcriptional response to fasting and refeeding in mice. Cells 8, 1529. 10.3390/cells8121529 (2019).

25 Brooks, R. C. & Garratt, M. G. Life history evolution, reproduction, and the origins of sex_dependent aging and longevity. Annals of the New York Academy of Sciences 1389, 92–107. 10.1111/nyas.13302 (2017).

26 Parihar, M. et al. Sex-dependent lifespan extension of ApcMin/+ FAP mice by chronic mTOR inhibition. Aging pathobiology and therapeutics 2, 187. 10.31491/apt.2020.12.039 (2020).

27 Bronikowski, A. M. et al. Sex_specific aging in animals: perspective and future directions. Aging cell 21, e13542. 10.1111/acel.13542 (2022).

28 Mitchell, S. J. et al. Effects of sex, strain, and energy intake on hallmarks of aging in mice. Cell metabolism 23, 1093–1112. 10.1016/j.cmet.2016.05.027 (2016).

29 Magwere, T., Chapman, T. & Partridge, L. Sex differences in the effect of dietary restriction on life span and mortality rates in female and male Drosophila melanogaster. The Journals of Gerontology Series A: Biological Sciences and Medical Sciences 59, B3–B9. 10.1093/gerona/59.1.B3 (2004).

30 Camus, M. F., Piper, M. D. & Reuter, M. Sex-specific transcriptomic responses to changes in the nutritional environment. Elife 8, e47262. 10.7554/eLife.47262 (2019).

31 Maklakov, A. A. et al. Sex-specific fitness effects of nutrient intake on reproduction and lifespan. Current Biology 18, 1062–1066. 10.1016/j.cub.2008.06.059 (2008).

32 Reda, G. K. et al. Dietary restriction and life-history trade-offs: insights into mTOR pathway regulation and reproductive investment in Japanese quails. bioRxiv 2023.11.14.567012. 10.1101/2023.11.14.567012 (2023).

33 Baloni, P. et al. Genome-scale metabolic model of the rat liver predicts effects of diet restriction. Sci Rep 9, 9807. 10.1038/s41598-019-46245-1 (2019).

34 National Research Council. Nutrient Requirements of Poultry. 9th Edition edn, (National Academy Press, 1994).

35 Meng, H., Yang, Y., Gao, Z.-H. & Wei, J.-H. Selection and validation of reference genes for gene expression studies by RT-PCR in Dalbergia odorifera. Scientific Reports 9, 3341. 10.1038/s41598-019-39088-3 (2019).

36 Pfaffl, M. W. Quantification strategies in real-time PCR. AZ of quantitative PCR 1, 89–113 2004).

37 Livak, K. J. & Schmittgen, T. D. Analysis of relative gene expression data using real-time quantitative PCR and the 2−ΔΔCT method. Methods 25, 402–408. 10.1006/meth.2001.1262 (2001).

38 R Core Team. R: A language and environment for statistical computing (Vienna, Austria, 2021). https://R-project.org/.

39 Wickham, H. ggplot2: Elegant Graphics for Data Analysis (Springer-Verlag New York, 2016). https://ggplot2.tidyverse.org

40 Burnham, K. P. & Anderson, D. R. Model selection and multimodel inference: a practical information-theoretic approach. (Springer, 2010).

41 Harrison, X. A. et al. A brief introduction to mixed effects modelling and multi-model inference in ecology. PeerJ 6, e4794. 10.7717/peerj.4794 (2018).

42 Bates, D., Mächler, M., Bolker, B. & Walker, S. Fitting Linear Mixed-Effects Models Using lme4. Journal of Statistical Software 67, 1–48. 10.18637/jss.v067.i01 (2015).

43 Kuznetsova, A., Brockhoff, P. B. & Christensen, R. H. B. LmerTest package: Tests in linear mixed effects models. Journal of Statistical Software 82, 1–26. 10.18637/jss.v082.i13 (2017).

44 Lenth, R., Singmann, H., Love, J., Buerkner, P. & Herve, M. (2018).

45 Vu, V. Q. ggbiplot: A ggplot2 based biplot (2011). http://github.com/vqv/ggbiplot

46 Kaiser, H. F. The application of electronic computers to factor analysis. Educational and psychological measurement 20, 141–151. 10.1177/00131644600200 (1960).

47 Chen, E.-H., Hou, Q.-L., Wei, D.-D., Jiang, H.-B. & Wang, J.-J. Phenotypic plasticity, trade-offs and gene expression changes accompanying dietary restriction and switches in Bactrocera dorsalis (Hendel)(Diptera: Tephritidae). Sci Rep 7, 1988. 10.1038/s41598-017-02106-3 (2017).

48 Raubenheimer, D., Hou, R., Dong, Y., Ren, C. & Cui, Z. Towards an integrated understanding of dietary phenotypes. Philosophical Transactions of the Royal Society B 378, 20220545. 10.1098/rstb.2022.0545 (2023).

49 Kilberg, M., Pan, Y.-X., Chen, H. & Leung-Pineda, V. Nutritional control of gene expression: how mammalian cells respond to amino acid limitation. Annual Review of Nutrition 25, 59. 10.1146/annurev.nutr.24.012003.132145 (2005).

50 Clarke, S. D. & Abraham, S. Gene expression: nutrient control of pre_and posttranscriptional events 1. The FASEB journal 6, 3146–3152. 10.1096/fasebj.6.13.1397836 (1992).

51 Feige-Diller, J. et al. The Impact of Varying Food Availability on Gene Expression in the Liver: Testing the Match-Mismatch Hypothesis. Frontiers in Nutrition 9. 10.3389/fnut.2022.910762 (2022).

52 de Sousa Abreu, R., Penalva, L. O., Marcotte, E. M. & Vogel, C. Global signatures of protein and mRNA expression levels. Molecular BioSystems 5, 1512–1526 2009).

53 Buccitelli, C. & Selbach, M. mRNAs, proteins and the emerging principles of gene expression control. Nature Reviews Genetics 21, 630–644. 10.1038/s41576-020-0258-4 (2020).

54 Chi, Y. et al. Regulation of gene expression during the fasting-feeding cycle of the liver displays mouse strain specificity. Journal of Biological Chemistry 295, 4809–4821. 10.1074/jbc.RA119.012349 (2020).

55 Gatti, D. et al. Genome_level analysis of genetic regulation of liver gene expression networks. Hepatology 46, 548–557. 10.1002/hep.21682 (2007).

56 Gokarn, R. et al. Long-term dietary macronutrients and hepatic gene expression in aging mice. J Gerontol A Biol Sci Med Sci 73, 1618–1625. 10.1093/gerona/gly065 (2018).

57 Horváthová, T., Nakagawa, S. & Uller, T. Strategic female reproductive investment in response to male attractiveness in birds. Proceedings of the Royal Society B 279, 163–170. 10.1098/rspb.2011.0663 (2012).

58 Marn, N., et al. Energetic basis for bird ontogeny and egg-laying applied to the bobwhite quail. Conservation Physiology 10, coac063. 10.1093/conphys/coac063 (2022).

59 Butler, A. A. & LeRoith, D. Minireview: tissue-specific versus generalized gene targeting of the igf1 and igf1r genes and their roles in insulin-like growth factor physiology. Endocrinology 142, 1685–1688. 10.1210/endo.142.5.8148 (2001).

60 Forbes, B. E., Blyth, A. J. & Wit, J. M. Disorders of IGFs and IGF-1R signaling pathways. Molecular and Cellular Endocrinology 518, 111035. 10.1016/j.mce.2020.111035 (2020).

61 Lodjak, J., Mänd, R. & Mägi, M. Insulin_like growth factor 1 and life_history evolution of passerine birds. Functional Ecology 32, 313–323. 10.1111/1365-2435.12993 (2018).

62 Ipsa, E., Cruzat, V. F., Kagize, J. N., Yovich, J. L. & Keane, K. N. Growth hormone and insulin-like growth factor action in reproductive tissues. Frontiers in endocrinology 10, 777. 10.3389/fendo.2019.00777 (2019).

63 Neirijnck, Y., Papaioannou, M. D. & Nef, S. The insulin/IGF system in mammalian sexual development and reproduction. International journal of molecular sciences 20, 4440. 10.3390/ijms20184440 (2019).

64 Tóth, Z., Mahr, K., Ölveczki, G., Őri, L. & Lendvai, Á. Z. Food restriction reveals individual differences in insulin-like growth factor-1 reaction norms. Frontiers in Ecology and Evolution 10, 826968. 10.3389/fevo.2022.826968 (2022).

65 Ashpole, N. M. et al. IGF-1 has sexually dimorphic, pleiotropic, and time-dependent effects on healthspan, pathology, and lifespan. Geroscience 39, 129–145. 10.1007/s11357-017-9971-0 (2017).

66 Meter, B., Kratochvíl, L., Kubička, L. & Starostová, Z. Development of male-larger sexual size dimorphism in a lizard: IGF1 peak long after sexual maturity overlaps with pronounced growth in males. Frontiers in Physiology 13, 917460. 10.3389/fphys.2022.917460 (2022).

67 Tóth, Z., Ouyang, J. Q. & Lendvai, Á. Z. Exploring the mechanistic link between corticosterone and insulin-like growth factor-1 in a wild passerine bird. PeerJ 6, e5936. 10.7717/peerj.5936 (2018).

68 McMurtry, J., Francis, G. & Upton, Z. Insulin-like growth factors in poultry. Domestic Animal Endocrinology 14, 199–229. 10.1016/S0739-7240(97)00019-2 (1997).

69 Bacon, W., Nestor, K., Emmerson, D., Vasilatos-Younken, R. & Long, D. Circulating IGF-I in plasma of growing male and female turkeys of medium and heavy weight lines. Domestic Animal Endocrinology 10, 267–277. 10.1016/0739-7240(93)90031-6 (1993).

70 Burnside, J. & Cogburn, L. A. Developmental expression of hepatic growth hormone receptor and insulin-like growth factor-I mRNA in the chicken. Molecular and Cellular Endocrinology 89, 91–96. 10.1016/0303-7207(92)90214-Q (1992).

71 Kita, K. et al. Influence of nutrition on hepatic IGF-I mRNA levels and plasma concentrations of IGF-I and IGF-II in meat-type chickens. Journal of Endocrinology 149, 181–190. 10.1677/joe.0.1490181 (1996).

72 Kita, K. Refeeding increases hepatic insulin-like growth factor-I (IGF-I) gene expression and plasma IGF-I concentration in fasted chicks. British poultry science 39, 679–682. 10.1080/00071669888566 (1998).

73 Giachetto, P. F. et al. Hepatic mRNA expression and plasma levels of insulin-like growth factor-I (IGF-I) in broiler chickens selected for different growth rates. Genetics and Molecular Biology 27, 39–44. 10.1590/s1415-47572004000100007 (2004).

74 Dauncey, M. et al. Nutritional regulation of growth hormone receptor gene expression. The FASEB journal 8, 81–88. 10.1096/fasebj.8.1.7507871 (1994).

75 Maes, M., Maiter, D., Ketelslegers, J.-M., Thissen, B. J.-P. & Underwood, L. Contributions of growth hormone receptor and postreceptor defects to growth hormone resistance in malnutrition. Trends in Endocrinology & Metabolism 2, 92–97. 10.1016/S1043-2760(05)80003-7 (1991).

76 Wang, Y. et al. Reduced serum insulin-like growth factor (IGF) I is associated with reduced liver IGF-I mRNA and liver growth hormone receptor mRNA in food-deprived cattle. The Journal of nutrition 133, 2555–2560. 10.1093/jn/133.8.2555 (2003).

77 Walock, C. N., Kittilson, J. D. & Sheridan, M. A. Characterization of a novel growth hormone receptor-encoding cDNA in rainbow trout and regulation of its expression by nutritional state. Gene 533, 286–294. 10.1016/j.gene.2013.09.046 (2014).

78 Deng, L., Zhang, W., Lin, H. & Cheng, C. H. Effects of food deprivation on expression of growth hormone receptor and proximate composition in liver of black seabream Acanthopagrus schlegeli. Comp Biochem Physiol 137, 421–432. 10.1016/j.cbpc.2004.01.008 (2004).

79 Li, Y. et al. Effect of early feed restriction on myofibre types and expression of growth-related genes in the gastrocnemius muscle of crossbred broiler chickens. British Journal of Nutrition 98, 310–319. 10.1017/S0007114507699383 (2007).

80 Swindell, W. R. Genes and gene expression modules associated with caloric restriction and aging in the laboratory mouse. BMC Genomics 10, 1–28. 10.1186/1471-2164-10-585 (2009).

81 Ham, D. J. et al. Distinct and additive effects of calorie restriction and rapamycin in aging skeletal muscle. Nature communications 13, 1–20. 10.1038/s41467-022-29714-6 (2022).

82 Yu, J. & Henske, E. P. Estrogen-induced activation of mammalian target of rapamycin is mediated via tuberin and the small GTPase Ras homologue enriched in brain. Cancer Research 66, 9461–9466. 10.1158/0008-5472.CAN-06-1895 (2006).

83 Piekarski, A. et al. Tissue distribution, gender-and genotype-dependent expression of autophagy-related genes in avian species. PLoS One 9, e112449. 10.1371/journal.pone.0112449 (2014).

84 Karanasios, E. et al. Autophagy initiation by ULK complex assembly on ER tubulovesicular regions marked by ATG9 vesicles. Nature Communications 7, 12420. 10.1038/ncomms12420 (2016).

85 Judith, D. et al. ATG9A shapes the forming autophagosome through Arfaptin 2 and phosphatidylinositol 4-kinase IIIβ. Journal of Cell Biology 218, 1634–1652. 10.1083/jcb.201901115 (2019).

86 Yang, C., Xia, S., Zhang, W., Shen, H.-M. & Wang, J. Modulation of Atg genes expression in aged rat liver, brain, and kidney by caloric restriction analysed via single-nucleus/cell RNA sequencing. Autophagy, 1–10. 10.1080/15548627.2022.2091903 (2022).

87 Green, C. L., Lamming, D. W. & Fontana, L. Molecular mechanisms of dietary restriction promoting health and longevity. Nature Reviews Molecular Cell Biology 23, 56–73. 10.1038/s41580-021-00411-4 (2022).

88 Jia, K. & Levine, B. Autophagy is required for dietary restriction-mediated life span extension in C. elegans. Autophagy 3, 597–599. 10.4161/auto.4989 (2007).

89 Morselli, E. et al. Caloric restriction and resveratrol promote longevity through the Sirtuin-1-dependent induction of autophagy. Cell death & disease 1, e10–e10. 10.1038/cddis.2009.8 (2010).

90 Martina, J. A., Chen, Y., Gucek, M. & Puertollano, R. mTORC1 functions as a transcriptional regulator of autophagy by preventing nuclear transport of TFEB. Autophagy 8, 903–914. 10.4161/auto.19653 (2012).

91 Sciarretta, S., Forte, M., Frati, G. & Sadoshima, J. The complex network of mTOR signalling in the heart. Cardiovascular Research 118, 424–439. 10.1093/cvr/cvab033 (2022).

92 Um, S. H., D’Alessio, D. & Thomas, G. Nutrient overload, insulin resistance, and ribosomal protein S6 kinase 1, S6K1. Cell Metabolism 3, 393–402. 10.1016/j.cmet.2006.05.003 (2006).

93 Ma, L. et al. Effect of caloric restriction on the SIRT1/mTOR signaling pathways in senile mice. Brain Research Bulletin 116, 67–72. 10.1016/j.brainresbull.2015.06.004 (2015).

