## Supplementary information for "Dietary restriction reveals sex-specific expression of the mTOR pathway genes in Japanese quails"

### Supplementary Tables

**Supplementary Table S1**. Ingredient composition and nutrient level of standardised basal feed for experimental quails

| Feed ingredients | Inclusion rate, % |
|---|---|
| Corn | 30.37 |
| Wheat | 20 |
| Soybean meal (46% CP) | 34.88 |
| Sunflower oil | 6.79 |
| Limestone | 5.64 |
| MCP | 1.29 |
| Salt | 0.38 |
| DL-Methionine | 0.15 |
| Vitamin and mineral premix[a] | 0.5 |
| **Nutrient content** | |
| Metabolisable energy MJ/kg | 12.13 |
| Crude protein | 20.0 |
| Calcium | 2.5 |
| Available Phosphorus | 0.35 |
| Sodium | 0.15 |
| Methionine | 0.45 |
| Methionine + cysteine | 0.75 |
| Lysine | 1.08 |
| Threonine | 0.74 |
| Leucine | 1.59 |
| Isoleucine | 0.86 |
| Arginine | 1.33 |
| Tryptophan | 0.25 |

[a] 1 kg premix provided: 1000000 NE vitamin A, 200 000 NE vitamin $D_3$, 4900 mg/kg vitamin E, 200 mg vitamin $K_3$, 150 mg vitamin $B_1$, 500 mg vitamin $B_2$, 1200 mg Ca-d-Pantothetane, 400 mg vitamin $B_6$, 2 mg vitamin $B_{12}$, 11 mg biotin, 2502 mg niacin, 60 mg folic acid, 300000 mg choline cloride, 13200 mg Zn, 1920 mg Cu, 9612 mg Fe, 13200 mg Mn, 180 mg I, 42 mg Se, 12 mg Co.

**Supplementary Table S2**. Average daily food intakes (ADFI (g)) of female and male groups during pre-treatment and week 1 and week 2 of the treatment period.

| | | Pre-treatment* | | Week 1 | | Week 2 | |
|---|---|---|---|---|---|---|---|
| Sex | Treatment | ADFI | SEM | ADFI | SEM | ADFI | SEM |
| | ADL | 30.41 | 1.07 | 33.119 | 1.153 | 32.784 | 1.821 |
| | DR20 | 29.70 | 1.11 | 23.275 | 0.761 | 23.568 | 0.891 |
| | DR30 | 30.08 | 1.41 | 20.156 | 1.554 | 20.178 | 1.547 |
| Female | DR40 | 29.76 | 0.51 | 17.855 | 0.304 | 17.537 | 0.299 |
| | ADL | 19.36 | 1.39 | 20.360 | 0.846 | 19.927 | 0.640 |
| | DR20 | 19.10 | 1.46 | 14.793 | 1.134 | 14.659 | 1.034 |
| | DR30 | 21.27 | 0.79 | 16.730 | 0.645 | 16.604 | 0.695 |
| Male | DR40 | 19.31 | 1.63 | 11.465 | 0.924 | 11.477 | 0.972 |

*the ADFI of DR20, DR30 and DR40 were calculated from the pre-treatment ADFI of individual birds. Abbreviations: ADFI, average daily feed intake; SEM, standard error of mean; ADL, *ad libitum*; DR20, 20% restriction; DR30, 30% restriction; DR40, 40% restriction

**Supplementary Table S3**. Characteristics of the primer pairs used. All amplifications were specific.

| Gene* | Primer sequences (5' → 3') | Annealing temp (°C) | NCBI GenBank | Product size (bp) | Primer efficiency | Observed Tm (°C) |
|---|---|---|---|---|---|---|
| *ACTB_F* | CCC CTG AAC CCC AAA GCC AAC | 56.2 | XM_015876619.1 | 114 | 109.47% | 84.9 |
| *ACTB_R* | ACC AGA GGC ATA CAG GGA CAG C | 56.1 | | | | |
| *GAPDH_F* | GCA CTG CGC CAC CTT CTC ACT | 57.5 | XM_015873412.2 | 116 | 104.95% | 87.7 |
| *GAPDH_R* | TGA CCA GGC GGC CAA TAC GG | 57.5 | | | | |
| *RN18s_F* | CCC TGC CGG AGC GTC GAG AA | 59.4 | XR_006936397.1 | 102 | 108.0% | 83.8 |
| *RN18s_R* | CCG GTA ATG ATC CTT CCG CAG GT | 56.1 | | | | |
| *mTOR_F* | CCG AAG CAT TGA ATT GGC CCT | 53.5 | XM_015882433.2 | 116 | 102.17% | 84.3 |
| *mTOR_R* | CAT CTC TCA AAG GCA GCG GAC C | 55.2 | | | | |
| *RPS6K1_F* | AGG CAG GAA CCC TCC GTG CAA | 58.8 | XM_015883670.2 | 106 | 106.05% | 85.6 |
| *RPS6K1_R* | AAG CTC AAA CTG CGA AGG GTC GG | 56.9 | | | | |
| *IGF1_F* | CAC TAT GCG GTG CTG AGC TGG TT | 55.8 | XM_015867574.2 | 118 | 105.5% | 84.3 |
| *IGF1_R* | ATC CCC TTG TGG TGT AAG CGT CT | 55.4 | | | | |
| *IGF1R_F* | TAC AAC TAC CGC TGC TGG ACC AC | 56.0 | XM_015873184.2 | 107 | 105.43% | 86.7 |
| *IGF1R_R* | AGG CAC TCA GGA TGG CAA CAC | 55.0 | | | | |
| *ATG9A_F* | CAA CGC CCT CAG GAT CCC CAT | 56.8 | XM_015868966.2 | 69 | 102.11% | 85.5 |
| *ATG9A_R* | ACG ATG CGG GCC TGT ACC TCC | 59.0 | | | | |
| *GHR_F* | GGC ACT GGT CTG TGT GAA TGA CT | 57.93 | XM_032441512.1 | 89 | 108.85% | 83.3 |
| *GHR_R* | CCA GCT CAG GTG ATC TGC ACT T | 57.58 | | | | |

* *ACTB*, beta-actin; *GAPDH*, glyceraldehyde-3-phosphate dehydrogenase; *RN18s*, 18S ribosomal RNA; *mTOR*, mechanistic target of rapamycin, *RPS6K1*, ribosomal protein S6 kinase 1, *IGF1*, insulin-like growth factor 1, *IGF1R*, insulin-like growth factor 1 receptor, *ATG9A*, autophagy-related 9A, *GHR*, growth hormone receptor.

**Supplementary Table S4**. Model Selection for male Japanese quails considering treatment, treatment period, bird identity, and experimental block as variables.

| Response variable | Model | AICc | df |
|---|---|---|---|
| **Body mass** | **~ treatment * week + (1\|birdID)** | **711.80** | **14** |
| Body mass | ~ treatment * week + (1\|block) | 743.45 | 14 |
| Body mass | ~ treatment * week + (1\|block/birdID) | 713.97 | 15 |

Abbreviations: DR, dietary restriction; week, restriction period; birdID, individual bird identity; block, experimental block; AICc, Akaike's information criterion corrected

**Supplementary Table S5**. Pairwise comparison of body mass of male Japanese quails in all dietary restriction levels at different time points.

| **Initial (day 0)** | | | | | |
|---|---|---|---|---|---|
| ***Contrast*** | ***Estimate*** | SEM | df | ***t-ratio*** | ***p-value*** |
| ADL – DR20 | 4.21 | 7.59 | 42.9 | 0.55 | 0.945 |
| ADL – DR30 | -5.45 | 7.59 | 42.9 | -0.72 | 0.889 |
| ADL – DR40 | 5.64 | 7.59 | 42.9 | 0.74 | 0.879 |
| DR20 – DR30 | -9.66 | 7.59 | 42.9 | -1.27 | 0.585 |
| DR20 – DR40 | 1.42 | 7.59 | 42.9 | 0.19 | 0.997 |
| DR30 – DR40 | 11.09 | 7.59 | 42.9 | 1.46 | 0.469 |
| **Week 1 (day 7)** | | | | | |
| ADL – DR20 | 15.53 | 7.59 | 42.9 | 2.05 | 0.187 |
| ADL – DR30 | 14.12 | 7.59 | 42.9 | 1.86 | 0.259 |
| ADL – DR40 | 21.9 | 7.59 | 42.9 | 2.89 | 0.030 |
| DR20 – DR30 | -1.40 | 7.59 | 42.9 | -0.18 | 0.997 |
| DR20 – DR40 | 6.38 | 7.59 | 42.9 | 0.84 | 0.835 |
| DR30 – DR40 | 7.78 | 7.59 | 42.9 | 1.02 | 0.736 |
| **Week 2 (day 14)** | | | | | |
| ADL – DR20 | 16.31 | 7.59 | 42.9 | 2.15 | 0.154 |
| ADL – DR30 | 15.01 | 7.59 | 42.9 | 1.98 | 0.212 |
| ADL – DR40 | 31.66 | 7.59 | 42.9 | 4.17 | <.001 |
| DR20 – DR30 | -1.30 | 7.59 | 42.9 | -0.17 | 998 |
| DR20 – DR40 | 15.35 | 7.59 | 42.9 | 2.02 | 0.196 |
| DR30 – DR40 | 16.65 | 7.59 | 42.9 | 2.19 | 0.141 |

We calculated mean body mass comparison of male birds among treatment groups within different time points using the function 'emmeans' with $p < 0.05$ significance level.

Abbreviations: ADL, *ad libitum*; DR20, 20% restriction; DR30, 30% restriction; DR40, 40% restriction. Initial, day 0; week 1, day 7; week 2, day 14.

**Supplementary Table S6**. Pairwise comparison of body mass of male Japanese quails in all the restriction time points at each restriction levels.

| | | | | | |
|---|---|---|---|---|---|
| | | ***Ad libitum group*** | | | |
| ***Contrast*** | ***Estimate*** | SEM | df | ***t-ratio*** | ***p-value*** |
| Initial – week 1 | -0.54 | 4.21 | 56 | -0.128 | 0.991 |
| Initial – week 2 | -0.16 | 4.21 | 56 | -0.039 | 0.999 |
| Week 1 – week 2 | 0.37 | 4.21 | 56 | 0.089 | 0.996 |
| | | **20% restricted group** | | | |
| Initial – week 1 | 10.77 | 4.21 | 56 | 2.56 | 0.035 |
| Initial – week 2 | 11.94 | 4.21 | 56 | 2.83 | 0.017 |
| Week 1 – week 2 | 1.16 | 4.21 | 56 | 0.28 | 0.959 |
| | | **30% restricted group** | | | |
| Initial – week 1 | 19.04 | 4.21 | 56 | 4.52 | <.001 |
| Initial – week 2 | 20.30 | 4.21 | 56 | 4.82 | <.001 |
| Week 1 – week 2 | 1.26 | 4.21 | 56 | 0.30 | 0.952 |
| | | **40% restricted group** | | | |
| Baseline – week 1 | 19.04 | 4.21 | 56 | 3.73 | 0.001 |
| Baseline – week 2 | 20.30 | 4.21 | 56 | 6.14 | <.001 |
| Week 1 – week 2 | 1.26 | 4.21 | 56 | 2.41 | 0.050 |

We analysed mean body mass comparison of male birds among time points within each treatment level using the function ‘emmeans’ with $p < 0.05$ significance level

**Supplementary Table S7**. Model Selection for sex-specific effects of treatments across time points considering sex, treatment and treatment period as fixed effects, and bird identity and experimental block as random variables.

| Response variable | Model | AICc | Df |
|---|---|---|---|
| **Body mass** | **~ treatment * sex * week + (1\|birdID)** | **1450.47** | **26** |
| Body mass | ~ treatment * sex * week + (1\|block) | 1522.94 | 26 |
| Body mass | ~ treatment * sex * week + (1\|block/birdID) | 1451.83 | 27 |

Abbreviations: DR, dietary restriction; week, restriction period; birdID, individual bird identity; block, experimental block; AICc, Akaike’s information criterion corrected

**Supplementary Table S8**. Pairwise comparison of body mass of females and males at all the dietary gradients and restriction period (week).

| **Treatment** | **Contrast** | **Time point** | **estimate** | ***SEM*** | ***df*** | ***t-ratio*** | ***p-value*** |
|---|---|---|---|---|---|---|---|
| ADL | Female – male | Initial mass | 43.25 | 8.42 | 85 | 5.14 | <0.001 |
| | Female – male | Week 1 mass | 51.66 | 8.42 | 85 | 6.14 | <0.001 |
| | Female – male | Week 2 mass | 49.17 | 8.42 | 85 | 5.84 | <0.001 |
| DR20 | Female – male | Initial mass | 44.01 | 8.42 | 85 | 5.23 | <0.001 |
| | Female – male | Week 1 mass | 29.10 | 8.42 | 85 | 3.46 | <0.001 |
| | Female – male | Week 2 mass | 22.75 | 8.42 | 85 | 2.70 | 0.008 |
| DR30 | Female – male | Initial mass | 24.98 | 8.42 | 85 | 2.97 | 0.004 |
| | Female – male | Week 1 mass | 17.99 | 8.42 | 85 | 2.14 | 0.035 |
| | Female – male | Week 2 mass | 7.06 | 8.42 | 85 | 0.84 | 0.404 |
| DR40 | Female – male | Initial mass | 34.01 | 8.42 | 85 | 4.04 | 0.001 |
| | Female – male | Week 1 mass | 11.22 | 8.42 | 85 | 1.33 | 0.186 |
| | Female – male | Week 2 mass | 9.82 | 8.42 | 85 | 1.18 | 0.246 |

We calculated mean body mass comparison between male and female birds within treatment groups and different time points using the function ‘emmeans’ with $p < 0.05$ significance level. Abbreviations: ADL, *ad libitum*; DR20, 20% restriction; DR30, 30% restriction; DR40, 40% restriction. Initial, day 0; week 1, day 7; week 2, day 14.

**Supplementary Table S9**. Pairwise comparison for the effect of dietary restriction groups on the expression of mTOR-related genes in male Japanese quails.

| **Genes** | **Contrast** | **Estimate** | SEM | *df* | ***t-ratio*** | ***p-value*** |
|---|---|---|---|---|---|---|
| mTOR | ADL – DR20 | 0.76 | 0.24 | 28 | 3.08 | 0.022 |
| | ADL – DR30 | 0.83 | 0.24 | 28 | 3.34 | 0.012 |
| | ADL – DR40 | 1.00 | 0.24 | 28 | 4.07 | 0.001 |
| | DR20 – DR30 | 0.06 | 0.24 | 28 | 0.26 | 0.994 |
| | DR20 – DR40 | 0.25 | 0.24 | 28 | 0.99 | 0.755 |
| | DR30 – DR40 | 0.18 | 0.24 | 28 | 0.74 | 0.812 |
| RPS6K1 | ADL – DR20 | -2.40 | 0.42 | 28 | -5.7 | <.001 |
| | ADL – DR30 | -2.37 | 0.42 | 28 | -5.66 | <.001 |
| | ADL – DR40 | -1.69 | 0.42 | 28 | -4.04 | 0.002 |
| | DR20 – DR30 | 0.02 | 0.42 | 28 | 0.06 | 0.999 |
| | DR20 – DR40 | 0.70 | 0.42 | 28 | 1.68 | 0.354 |
| | DR30 – DR40 | 0.67 | 0.42 | 28 | 1.62 | 0.384 |
| ATG9A | ADL – DR20 | -0.71 | 0.25 | 28 | -2.86 | 0.037 |
| | ADL – DR30 | -1.12 | 0.25 | 28 | -4.52 | <.001 |
| | ADL – DR40 | -0.78 | 0.25 | 28 | -3.18 | 0.018 |
| | DR20 – DR30 | -0.41 | 0.25 | 28 | -1.66 | 0.365 |
| | DR20 – DR40 | -0.07 | 0.25 | 28 | -0.30 | 0.990 |
| | DR30 – DR40 | 0.33 | 0.25 | 28 | 1.35 | 0.539 |
| IGF1 | ADL – DR20 | 1.94 | 0.41 | 28 | 4.75 | <.001 |
| | ADL – DR30 | 1.75 | 0.41 | 28 | 4.27 | <.001 |
| | ADL – DR40 | 2.65 | 0.41 | 28 | 6.47 | <.001 |
| | DR20 – DR30 | -0.19 | 0.41 | 28 | -0.47 | 0.964 |
| | DR20 – DR40 | 0.71 | 0.41 | 28 | 1.72 | 0.332 |
| | DR30 – DR40 | 0.90 | 0.41 | 28 | 2.19 | 0.149 |
| IGF1R | ADL – DR20 | 0.56 | 0.41 | 28 | 1.37 | 0.528 |
| | ADL – DR30 | 0.37 | 0.41 | 28 | 0.90 | 0.803 |
| | ADL – DR40 | 1.41 | 0.41 | 28 | 3.45 | 0.009 |
| | DR20 – DR30 | -0.19 | 0.41 | 28 | -0.47 | 0.965 |
| | DR20 – DR40 | 0.85 | 0.41 | 28 | 2.08 | 0.183 |
| | DR30 – DR40 | 1.04 | 0.41 | 28 | 2.55 | 0.074 |
| GHR | ADL – DR20 | -0.02 | 0.39 | 28 | 0.06 | 0.999 |
| | ADL – DR30 | -0.88 | 0.39 | 28 | -2.22 | 0.141 |
| | ADL – DR40 | -0.93 | 0.39 | 28 | -2.36 | 0.108 |
| | DR20 – DR30 | -0.90 | 0.39 | 28 | -2.28 | 0.126 |
| | DR20 – DR40 | -0.95 | 0.39 | 28 | -2.42 | 0.096 |
| | DR30 – DR40 | -0.05 | 0.39 | 28 | -0.14 | 0.999 |

We calculated mean gene expression comparison of male birds among treatment groups using the function 'emmeans' with $p < 0.05$ significance level. Abbreviations: *mTOR*, mechanistic target of rapamycin; *RPS6K1*, ribosomal protein S6 kinase 1; *ATG9A*, autophagy-related 9A; *IGF1*, insulin-like growth factor 1; *IGF1R*, insulin-like growth factor 1 receptor; *GHR*, growth hormone receptor; ADL, *ad libitum*; DR20, 20% restriction; DR30, 30% restriction; DR40, 40% restriction;

**Supplementary Table S10**. Pairwise comparison of expression of genes between females and males at all dietary levels.

| Treatment | Contrast | Time point | estimate | *SEM* | *df* | *t-ratio* | *p-value* |
|---|---|---|---|---|---|---|---|
| mTOR | ADL | Female – male | 0.91 | 0.30 | 56 | 2.99 | 0.004 |
| | DR20 | Female – male | 0.74 | 0.30 | 56 | 2.44 | 0.018 |
| | DR30 | Female – male | -0.06 | 0.30 | 56 | -0.21 | 0.831 |
| | DR40 | Female – male | -0.22 | 0.30 | 56 | -0.74 | 0.464 |
| RPS6K1 | ADL | Female – male | 0.65 | 0.36 | 56 | 1.78 | 0.080 |
| | DR20 | Female – male | -1.12 | 0.36 | 56 | -3.08 | 0.003 |
| | DR30 | Female – male | -0.64 | 0.36 | 56 | -1.74 | 0.087 |
| | DR40 | Female – male | 0.19 | 0.36 | 56 | 0.52 | 0.607 |
| ATG9A | ADL | Female – male | -0.26 | 0.25 | 56 | -1.04 | 0.304 |
| | DR20 | Female – male | -0.43 | 0.25 | 56 | -2.92 | 0.005 |
| | DR30 | Female – male | -0.11 | 0.25 | 56 | -4.48 | <.001 |
| | DR40 | Female – male | -0.21 | 0.25 | 56 | -0.85 | 0.395 |
| IGF1 | ADL | Female – male | 1.53 | 0.49 | 56 | 3.12 | 0.003 |
| | DR20 | Female – male | 1.14 | 0.49 | 56 | 2.32 | 0.024 |
| | DR30 | Female – male | 0.62 | 0.49 | 56 | 1.27 | 0.209 |
| | DR40 | Female – male | 2.43 | 0.49 | 56 | 4.96 | <.001 |
| IGF1R | ADL | Female – male | -0.26 | 0.37 | 56 | -0.78 | 0.476 |
| | DR20 | Female – male | -0.23 | 0.37 | 56 | -0.62 | 0.538 |
| | DR30 | Female – male | -0.35 | 0.37 | 56 | -0.95 | 0.347 |
| | DR40 | Female – male | 0.74 | 0.37 | 56 | 2.02 | 0.048 |
| GHR | ADL | Female – male | -0.29 | 0.44 | 56 | -0.67 | 0.505 |
| | DR20 | Female – male | -0.03 | 0.44 | 56 | 0.07 | 0.942 |
| | DR30 | Female – male | 0.62 | 0.44 | 56 | 1.41 | 0.164 |
| | DR40 | Female – male | 0.49 | 0.44 | 56 | 1.11 | 0.270 |

We calculated mean gene expression comparison between male and female birds within treatment groups using the function ‘emmeans’ with $p < 0.05$ significance level. Abbreviations: *mTOR*, mechanistic target of rapamycin; *RPS6K1*, ribosomal protein S6 kinase 1; *ATG9A*, autophagy-related 9A; *IGF1*, insulin-like growth factor 1; *IGF1R*, insulin-like growth factor 1 receptor; *GHR*, growth hormone receptor; ADL, *ad libitum*; DR20, 20% restriction; DR30, 30% restriction; DR40, 40% restriction.

**Supplementary Table S11**. Eigenvalues and respective complained variance of PCs from gene expression variables.

| PCs | Eigenvalue | Variance % | Cumulative variance % |
|---|---|---|---|
| PC1 | 2.5277708 | 42.129514 | 42.12951 |
| *PC1* | 1.6129661 | 26.882768 | 69.01228 |
| *PC3* | 0.6831661 | 11.386102 | 80.39838 |
| *PC4* | 0.5357689 | 8.929482 | 89.32786 |
| *PC5* | 0.3674151 | 6.123586 | 95.45145 |
| *PC6* | 0.2729130 | 4.548550 | 100.00 |

**Supplementary Table S12**. Contribution of original variables to the principal components.

| | ***Principal components*** | | | | | |
|---|---|---|---|---|---|---|
| ***Original variables*** | ***PC1*** | ***PC2*** | ***PC3*** | ***PC4*** | ***PC5*** | ***PC6*** |
| *mTOR* | 0.4256 | -0.4256 | 0.2501 | -0.2045 | 0.6430 | -0.3459 |
| *RPS6K1* | -0.4691 | -0.3231 | -0.3922 | -0.1516 | 0.4797 | 0.5181 |
| *ATG9A* | -0.4643 | -0.2821 | 0.1461 | -0.6597 | -0.3412 | -0.3628 |
| *IGF1* | -0.3271 | -0.4263 | 0.6481 | 0.5133 | -0.0748 | 0.1481 |
| *IGF1R* | 0.4998 | -0.2841 | 0.1610 | -0.3533 | -0.3577 | 0.6250 |
| *GHR* | 0.1620 | -0.6101 | -0.5623 | 0.3338 | -0.3259 | -0.2599 |

Abbreviations: *mTOR*, mechanistic target of rapamycin; *RPS6K1*, ribosomal protein S6 kinase 1; *ATG9A*, autophagy-related 9A; *IGF1*, insulin-like growth factor 1; *IGF1R*, insulin-like growth factor 1 receptor; *GHR*, growth hormone receptor

### Supplementary Figures

**Supplementary Figure S1**. Melting curves for each target and reference genes primer dimers. Abbreviations: *ACTB*, beta-actin; *mTOR*, mechanistic target of rapamycin; *RPS6K1*, ribosomal protein S6 kinase 1; *IGF1*, insulin-like growth factor 1; *IGF1R*, insulin-like growth factor 1 receptor; *ATG9A*, autophagy-related 9A; *GHR*, growth hormone receptor; *GAPDH*, glyceraldehyde-3-phosphate dehydrogenase; *RN18s*: 18S ribosomal RNA.

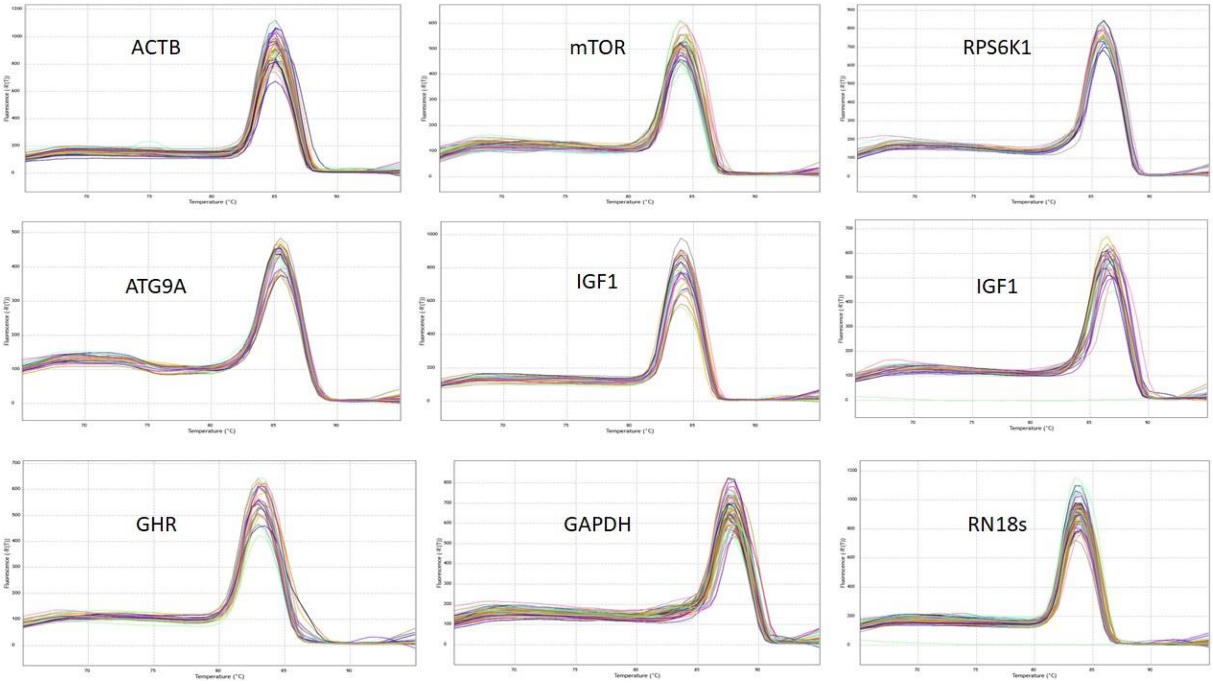

**Supplementary Figure S2**. Pearson correlation among expression of genes mediating nutrient availability. *mTOR*, mechanistic target of rapamycin; *RPS6K1*, ribosomal protein S6 kinase 1; *IGF1*, insulin-like growth factor 1; *IGF1R*, insulin-like growth factor 1 receptor; *ATG9A*, autophagy-related 9A; *GHR*, growth hormone receptor; *GAPDH*, glyceraldehyde-3-phosphate dehydrogenase; *RN18s*: 18S ribosomal RNA.

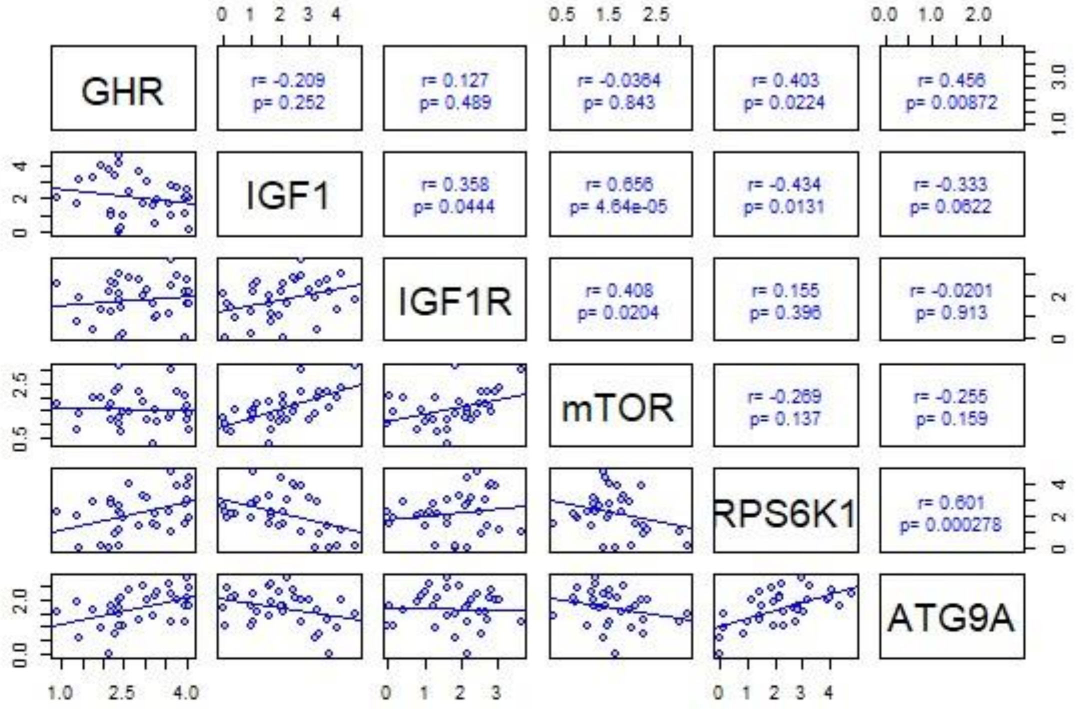


Most of the analysed genes were significantly related (figure S2). *IGF1* gene expression showed a significantly positive correlation with *IGF1R* and *mTOR*, whereas a negative correlation with *RPS6K1* and *ATG9A* gene expression. The *mTOR* also showed a positive correlation with *IGF1R*, while it was negatively correlated with the *RPS6K1* and *ATG9A* gene expression. The expression of *GHR*, *RPS6K1* and *ATG9A* genes were positively related. Unexpectedly, *GHR* expression had no significant correlation with the expression of *IGF1* and *mTOR* genes.
